# A failure to discriminate social from non-social touch at the circuit level may underlie social avoidance in autism

**DOI:** 10.1101/2024.06.19.599778

**Authors:** Trishala Chari, Ariana Hernandez, João Couto, Carlos Portera-Cailliau

**Affiliations:** Department of Neurology, David Geffen School of Medicine at the University of California Los Angeles Los Angeles, CA 90095; Neuroscience Interdepartmental Program, David Geffen School of Medicine at the University of California Los Angeles Los Angeles, CA 90095; Department of Neurobiology, David Geffen School of Medicine at the University of California Los Angeles Los Angeles, CA 90095

## Abstract

Social touch is critical for communication and to impart emotions and intentions. However, certain autistic individuals experience aversion to social touch, especially when it is unwanted. We used a novel social touch assay and Neuropixels probes to compare neural responses to social vs. non-social interactions in three relevant brain regions: vibrissal somatosensory cortex, tail of striatum, and basolateral amygdala. We find that wild type (WT) mice showed aversion to repeated presentations of an inanimate object but not of another mouse. Cortical neurons cared most about touch context (social vs. object) and showed a preference for social interactions, while striatal neurons changed their preference depending on whether mice could choose or not to interact. Amygdalar and striatal neurons were preferentially modulated by forced object touch, which was the most aversive. In contrast, the *Fmr1* knockout (KO) model of autism found social and non-social interactions equally aversive and displayed more aversive facial expressions to social touch when it invaded their personal space. Importantly, when *Fmr1* KO mice could choose to interact, neurons in all three regions did not discriminate social valence. Thus, a failure to differentially encode social from non-social stimuli at the circuit level may underlie social avoidance in autism.

## INTRODUCTION

The sense of touch is crucial to social communication and interaction, and animals constantly seek it, as manifested in humans by hugging, kissing, caressing, and even tickling one another. Through affiliative touch, animals offer comfort, provide inference about their internal states, and build or modify social relationships^1–5^. Brain circuits therefore likely evolved to prefer social touch over non-social stimuli^6,7^. On the other hand, social touch may be perceived as aversive when it is forced onto the subject^8^ or when it invades our peri-personal space^9^. Moreover, certain autistic individuals actively avoid social interactions^4,10,11^, perhaps because they are unable to discriminate the unique valence/salience of social stimuli, or to discriminate them from non-social stimuli.

The circuits involved in social touch are beginning to be elucidated^6,12^. In the somatosensory cortex, neural activity is modulated differently by social touch compared to non-social touch^13–16^. In rodent vibrissal somatosensory cortical (vS1), neurons are even capable of distinguishing between mice of different sexes^17^. This suggests that vS1 is not simply responding to shape or texture but is influenced by information about context coming from other regions known to be implicated in the encoding of social affiliative touch^18–20^.

Here we wanted to address three important questions related to how behavioral responses and neural activity are uniquely modulated by social touch: First, does activity in certain brain regions reflect an animal’s preference for social vs. non-social interactions? Second, do neural responses to social vs. non-social touch depend on whether the animal has a choice to interact (voluntary vs. forced touch)? This relates to the concept of personal space, the notion that touch might be more aversive when the individual has no option but to engage in it. And finally, do these circuits process social touch differently in individuals with neurodevelopmental conditions (NDCs), like autism spectrum disorder (ASD), who perceive social touch as aversive?

Using our recently developed social touch assay^21^, we first identified which brain areas are recruited by social touch and then used Neuropixels silicon probes^22^ to record neuronal responses to repeated interactions with either a stranger mouse or a plastic object. We focused on three relevant brain regions: vS1, because of its critical role in processing whisker-mediated touch^23^ and social touch in rats^15–17^; the basolateral amygdala (BLA) because of its involvement in emotional processing and the encoding of aversive and social stimuli^24–29^; and tail of the striatum (tSTR), which has been implicated in sensorimotor decision-making^30^ and in aversion to novelty in ASD mice^31^. In addition to wild type (WT) control mice, we examined the *Fmr1* knockout (KO) mouse model of Fragile X Syndrome (FXS), the most common single gene cause of ASD and intellectual disability^32^. *Fmr1* KO mice manifest tactile defensiveness to repetitive whisker stimulation^33,34^ and show greater avoidance/aversion to social touch that WT controls^21^.

We find that WT mice display significant aversion to object touch but not to social touch. In contrast, *Fmr1* KO mice fail to perceive the social valence by showing as much aversion to social and object touch and show unusually high aversion to social touch within their personal space. When mice can voluntary initiate touch, neurons in vS1 and tSTR of WT mice are preferentially modulated by social touch, whereas neurons in *Fmr1* KO mice are not. In contrast, under forced touch conditions, neurons in tSTR and BLA in both WT and KO mice responded most to forced object touch, which was the most aversive for both genotypes.

## RESULTS

### Wild type mice show avoidance and aversive facial expressions to object touch but not to social touch

We first asked if WT mice perceive social touch differently from non-social touch. We used the same behavioral assay we recently developed^21^ in which a head-fixed test mouse that can run on a polystyrene ball is exposed to repeated presentations of either an inanimate object (50 mL plastic tube) or a stranger mouse using a motorized platform (**Fig. 1a**). Each presentation bout lasted 10 s and consisted of a 5 s period when the platform was stopped at the touch position, and a 5 s interstimulus interval (ISI) during which the platform moved away from and back toward the test mouse (**Fig. 1a-b**; see Methods). To assess whether mice would respond differently depending on whether they could choose to engage with their visitor, presentations were either voluntary (the platform stops at a position in which the test mouse can willingly initiate contact with the visitor via its whiskers) or forced (the platform stops at a position where the snouts of the two mice are in direct physical contact). We acquired high-resolution videos and used FaceMap and DeepLabCut to quantify changes in facial expressions (aversive whisker protraction, orbital tightening) and in running direction (see Methods; **Fig. 1c**).

**Fig. 1:**
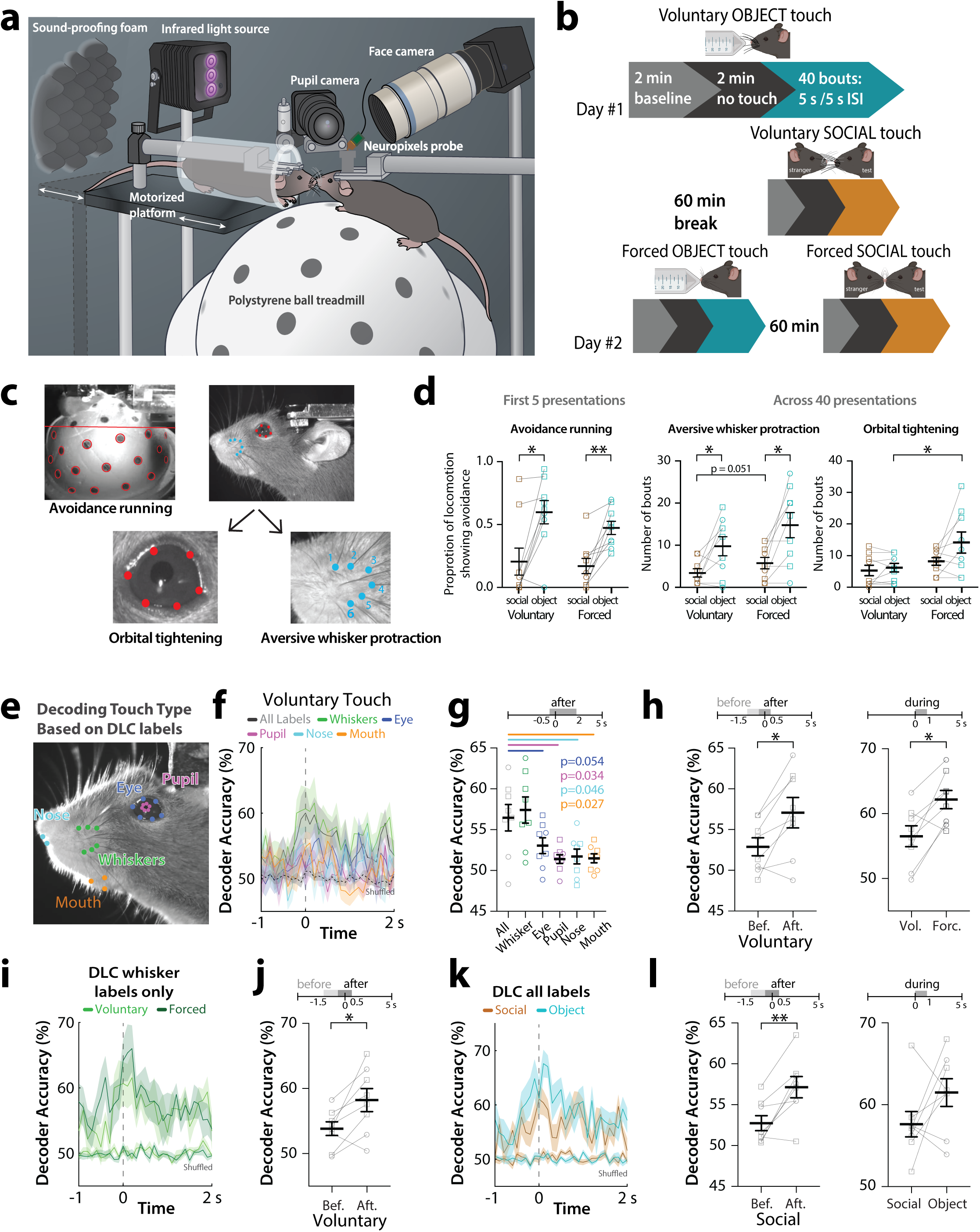
WT mice show avoidance and aversive facial expressions to object touch, but not social touch. **a.** Cartoon of behavioral assay for social facial touch. A head-fixed test mouse, chronically implanted with a Neuropixels 1.0 probe, can run on an air-suspended polystyrene ball while interacting with a stranger mouse restrained in a plexiglass tube secured to a motorized platform. The system is fully automated to move the stranger mouse to different distances away from the test mouse. Two cameras focus on the face and the eye/pupil, respectively, while a third camera that tracks the mouse and ball motion is overhead (not shown). An infrared light source provides light for tracking behavioral responses. Acoustic foam is used for sound insulation. **b.** Timeline of social touch behavioral assay. Mice receive 40 bouts of voluntary object and voluntary social touch, 60 min apart, on Day #1. On Day #2, mice receive 40 bouts of forced object and forced social touch. For each bout the platform is moving for 5 s and stopped for 5 s. **c.** Avoidance running and different aversive facial expressions (orbital tightening and aversive whisker protraction) are quantified using FaceMap and DeepLabCut. **d.** Running avoidance during the first 5 presentations of touch in WT mice (left). Number of bouts of prolonged whisker protraction (middle) and orbital tightening (right) across all 40 bouts of touch in WT mice. Squares=males, circles=females. **p<0.01, *p<0.05, two-way ANOVA with Bonferroni’s. **e.** The motion of DeepLabCut (DLC) labels corresponding to different facial features were used to decode touch context (social vs. object) and touch choice (voluntary vs. forced) with SVM classifiers. **f.** Decoder performance of touch context using DLC labels on the face in WT mice across time (from 1 s before to 2 s after platform stops). **g.** Decoder accuracy for context discrimination for using DLC labels for all facial features or labels for individual facial features (whiskers, eye, pupil, nose or mouth) during the time after platform movement (−0.5 s to +2 s) for voluntary touch. **h.** Left: Decoder accuracy for context discrimination using all DLC labels before (−1.5 to −0.5 s) and after (−0.5 to +0.5 s) platform movement. Right: Decoder accuracy for context discrimination using all DLC labels for voluntary touch and forced touch. p<0.01 for nonparametric Kruskal-Wallis test for left panel. *p<0.05 for parametric paired t-test for middle and right panel. **i.** Decoder performance for context discrimination using only DLC whisker labels in WT mice differently across time for voluntary and forced touch (from −1 to +2 s). **j.** Decoder accuracy for context discrimination using DLC whisker labels before (−1.5 to −0.5 s) and after (−0.5 to +0.5 s) platform movement for voluntary touch. **k.** Decoder performance of touch choice (voluntary vs, forced) using all DLC labels on the mouse’s face in WT mice across time for social and object touch (from −1 to +2 s). **l.** Decoder accuracy for choice discrimination using all DLC labels before (−1.5 to −0.5 s) and after (−0.5 to +0.5 s) platform movement for social touch (left) and during the first second for social versus object touch (right). **p<0.01 for parametric paired t-test for both panels.

We previously reported that WT mice display running avoidance and aversive facial expressions (AFEs) to forced object touch, but not nearly as much to forced social touch^21^. We once again observed that WT mice (n=9) spend a significantly greater proportion of their running time showing avoidance to object touch compared to social touch (**Fig. 1d**; voluntary p=0.012, forced p=0.005). WT mice also displayed significantly more bouts of aversive whisker protraction with object touch than with social touch across all 40 presentations (**Fig. 1d**; voluntary: p=0.049, forced: p=0.021), and a higher proportion of time displaying aversive whisker protraction with object touch (**Supplementary Fig. 1**; voluntary: p=0.075, forced: p=0.013). Thus, WT mice prefer social to non-social interactions, in line with our previous findings^21^.

We next examined whether responses are different when the animal can voluntarily initiate contact compared to when they have no choice. We found that forced touch elicited more aversive whisker protraction and orbital tightening (**Fig. 1d**; p=0.051 and p=0.026, respectively). Thus, unsolicited object touch is particularly aversive to WT mice.

### Touch context (social vs. object) can be decoded from facial expressions

Based on the above behavioral results, we hypothesized that it would be possible to accurately decode touch context from behavior videos. Hence, we trained a support vector machine (SVM) classifier on DeepLabCut (DLC) labels of orofacial movements (see Methods; **Fig. 1e**), which reflect how mice engage with their environment^35–37^. We found that whiskers contributed most to decoding accuracy of touch context (social vs. object), such that the performance of the whisker-based decoder was similar to that of the all-labels decoder (**Fig. 1f-g**; p>0.05). Interestingly, during voluntary touch presentations, the all-DLC decoder performance rose suddenly just before the platform stopped, as whiskers first made contact (**Fig. 1h**; **left**; before vs. after p=0.013). Decoding accuracy for touch context was also higher for forced touch compared to voluntary (**Fig. 1h, right**; p=0.012). When we trained the SVM classifier only on whisker DLC labels we also found better decoding accuracy after whisker contact (**Fig. 1i, j**; p=0.018).

A classifier trained to discriminate touch choice (voluntary vs. forced) also showed greater decoding accuracy upon movement of the platform towards the test mouse (**Fig. 1k,l**; p=0.005). Furthermore, we found a non-significant trend toward greater accuracy for decoding touch choice during object touch compared to social touch, which likely reflects the special aversion of WT mice to object touch when it is forced upon them within their personal space (**Fig. 1l**; **right**).

### Social touch engages neurons in cortical, striatal, amygdalar circuits

Circuits involved in social behaviors are widespread throughout the brain^12,38^. We sought to survey which brain regions are uniquely modulated by social touch. We used transgenic TRAP2 mice (cFos-Cre^ERT^^2^ x Ai14) in which the expression of tdTom is driven in an activity-dependent manner via the cFos promoter^39,40^. TRAP2 mice received repetitive presentations of either forced object (n=6) or social touch (n=6) for 30 min following induction with 4-hydroxytamoxifen (4-OHT) and were perfused 72 h later to quantify tdTom expression (**Fig. 2a**; see Methods). Control TRAP2 mice (n=5) were induced in a no-touch condition.

**Fig. 2:**
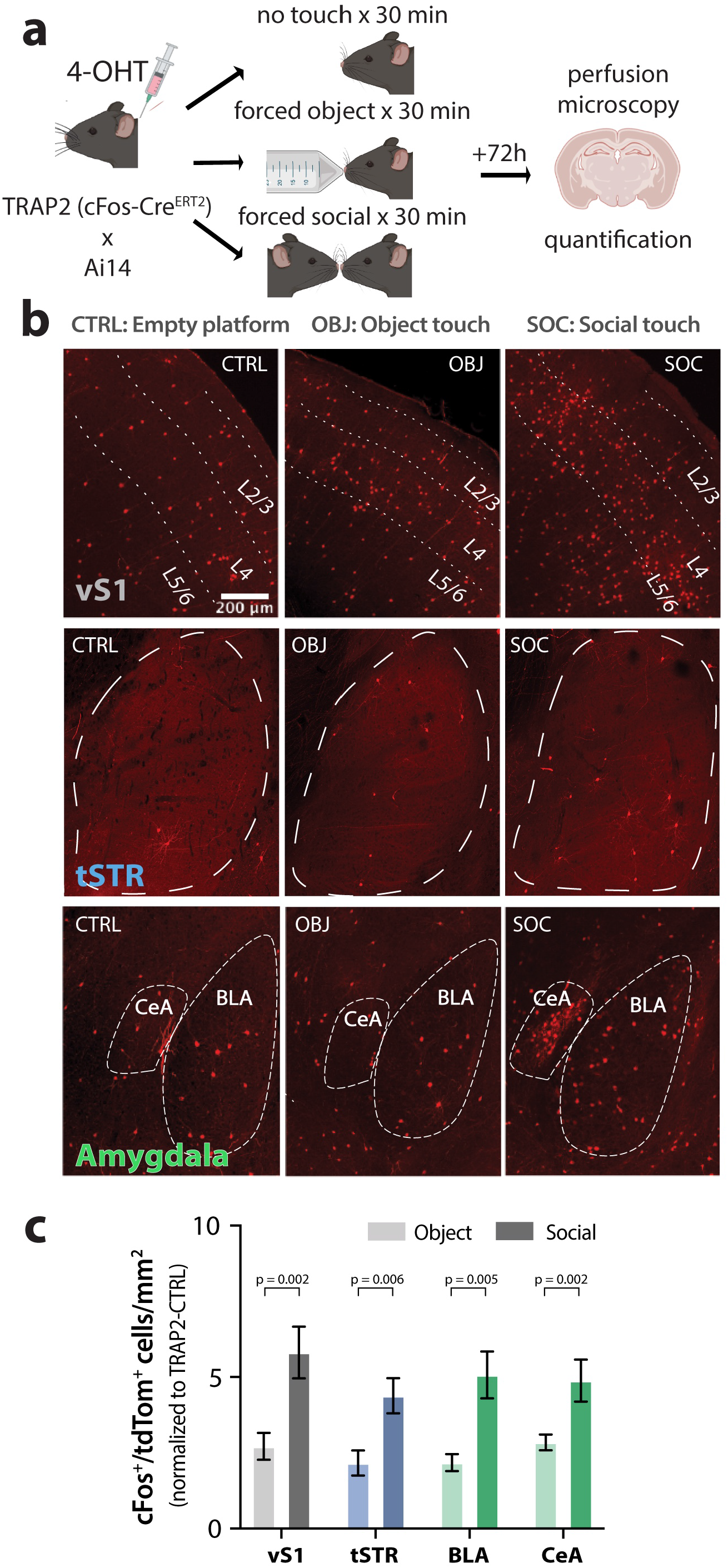
Differential cFos expression by forced object and social touch across vS1, tSTR and BLA of TRAP2 mice. **a.** Experimental protocol for TRAP2 behavioral experiments. TRAP2 WT mice were injected with 4-OHT 30 min prior to behavioral testing, and then received either repetitive bouts of social or object touch (5 s stim, 5 s ISI) for 30 min. As a control, a subset of mice received repetitive bouts of the same duration with the platform moving but without an object or mouse present (TRAP2-CTRL). Mice were perfused 72 h later and their brains sectioned for histological analysis and quantification of cFos expression. **b.** Example images of cFos expression in vS1, tSTR, and central amygdala (CeA) and BLA after object touch (OBJ), social touch (SOC), or no touch (empty platform; CTRL) (scale bar = 200 µm). **c.** Density of cFos-expressing (tdTom^+^) cells per mm^2^ for each brain region, normalized against CTRL density. *p<0.05, normality was tested with D’Agostino & Pearson test followed by unpaired nonparametric Mann-Whitney or parametric t-test for each brain region. Each bar represents data from 5-6 mice, and at least 6 images were collected from a single mouse for each brain region.

Forced object and social touch induced cFos expression across many regions throughout the brain. As expected, we identified *cFos* induction in layer 2/3 of the vS1 (**Fig. 2b,c**). Additionally, we observed high expression of tdTom in regions associated with social behavior and/or aversion, including the nucleus accumbens (NAc), the medial and basolateral amygdala (MeA, BLA), the paraventricular nucleus of the thalamus (PVT), the periaqueductal gray (PAG), and the insular cortex (ICx) (**Supplementary Fig. 2**). Importantly, tdTom expression was significantly higher after social touch than after object touch in L2/3 of vS1, the tSTR, the BLA, and the central amygdala (CeA) (**Fig. 2b,c**; vS1 p=0.002, tSTR p=0.006, BLA p=0.005, CeA p=0.002), as well as in anterior cingulate cortex (ACCx), ICx, BLA, MeA, PVT and the paraventricular nucleus of the hypothalamus (PVHy) (**Supplementary Fig. 2b**; ACCx p<0.001, NAc p=0.051, ICx p=0.008, BLA p=0.005, MeA p=0.030, PVT p=0.002, PVHy p=0.055). These cortical and subcortical brain regions are all known to be involved in social behavior and aversive processing^20,26,29–31,41–46^. In contrast, we did not observe significant differences between social and object touch in the density of tdTom-expressing cells in the PAG, primary motor cortex (MC), or visual cortex (V1) (**Supplementary Fig. 2b**). Thus, social touch preferentially recruits neurons in a subset of brain regions.

### vS1, tSTR and BLA neurons are differentially modulated by object vs. social touch

Based on the above TRAP2 results, we chose to implant single Neuropixels probes such that their trajectory would allow us to record simultaneously from vS1, tSTR, and BLA. In this way, we could investigate how social facial touch is differentially represented from non-social touch within sensory (vS1) and emotional-related brain areas (BLA), as well as within a sensorimotor-related brain region (tSTR). Furthermore, these brains regions have been shown to be involved during social and aversive behaviors^15,27,31,41,42,47^.

We chronically implanted single-shank Neuropixels silicon probes in 9 WT mice (**Fig. 3a**) and confirmed targeting of all 3 regions through histological reconstruction of the probe tract (**Supplementary Fig. 3**; see Methods). We used these probe trajectories and electrophysiological landmarks to putatively assign units to vS1, tSTR, or BLA (see Methods; **Supplementary Fig. 4a-c**). Only manually curated, isolated single units were considered in the analysis (see Methods).

**Fig. 3:**
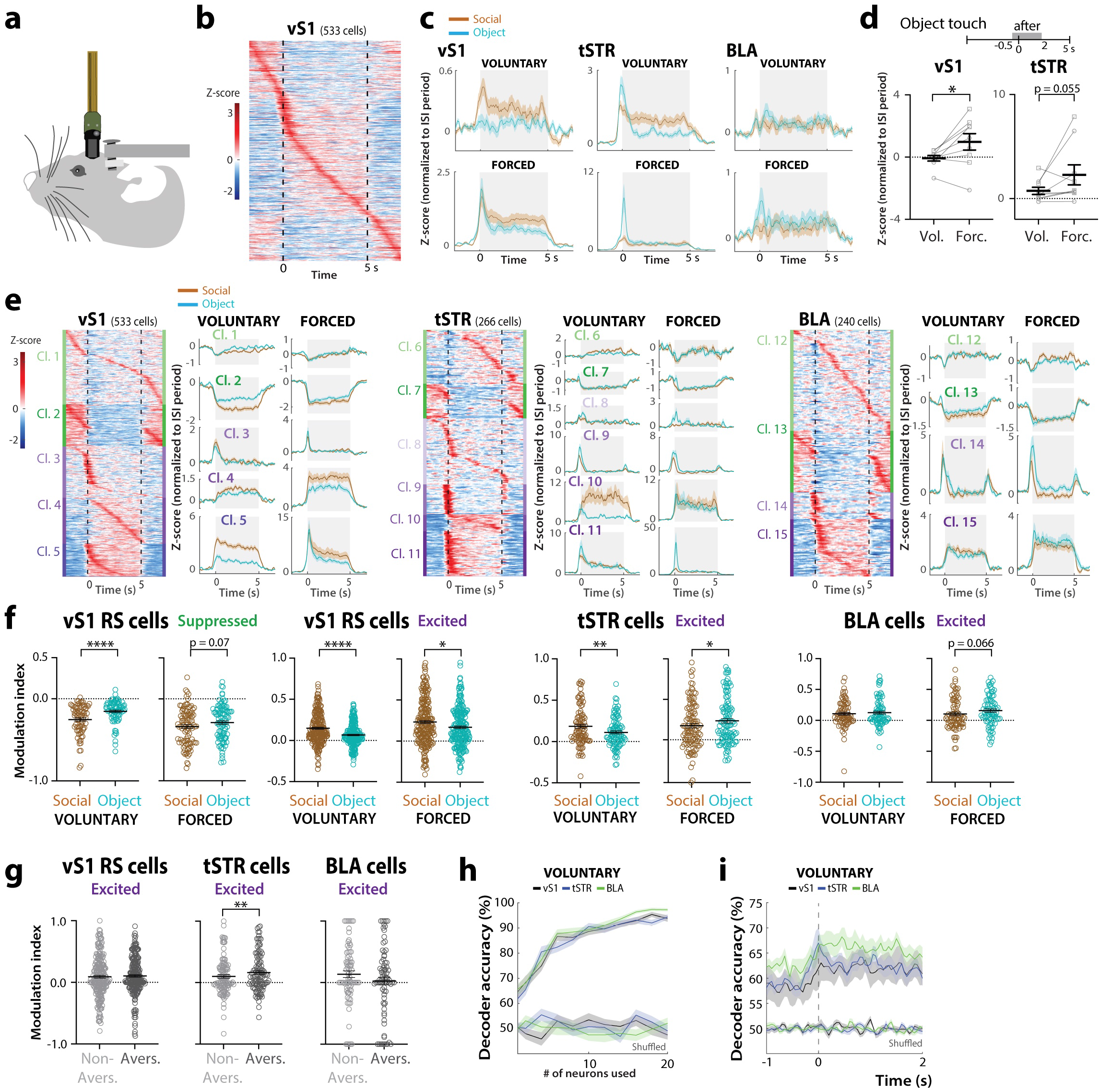
Neurons in vS1, tSTR and BLA show differential modulation by social vs. object touch. **a.** Cartoon of mouse chronically implanted with Neuropixels probe (not to scale). **b.** Example heatmap of all vS1 cells (n=533) sorted by peak of trial-averaged, z-scored PSTHs. Scale bar denotes z-score values. **c.** Trial-averaged z-scores normalized to the period before touch (ISI) for all vS1 regular-spiking (RS), tSTR and BLA cells from 9 WT mice during social touch (brown) and object touch (blue) for voluntary (top) and forced conditions (bottom) across time (−2 to +7 s). **d.** Mean z-scores during the period of contact (−0.5 to +2 s) for voluntary and forced object touch across different mice. *p<0.05 for paired parametric t-test. **e.** Left: Heatmap of the trial-averaged PSTHs for voluntary touch (taken as an average of all object and social touch presentations) for all vS1, tSTR and BLA cells split by clusters derived from PCA-k-means clustering and sorted by peak firing in time within each cluster. Right: Mean Z-scores for all neurons of individual clusters for social vs. object touch during voluntary or forced presentations, as a mean of all 40 presentations. Clusters are sorted by suppressed (green) to excited (purple) and shading indicates magnitude of how much they were suppressed/excited. Time 0 s denotes when the platform stops in the contact position. **f.** Modulation index of vS1 RS excited and suppressed cells, tSTR excited cells, and BLA excited cells for social vs. object touch. ****p<0.0001, **p<0.01, *p<0.05 for paired parametric t-test. Each symbol represents a single cell (n=9 WT mice). **g.** Modulation index of excited cells in vS1, tSTR, and BLA during forced object presentations when WT mice did not show aversion (Non-Avers.) or presentations when the same animals showed AFEs (Avers.). **h.** Decoder accuracy for touch context (social vs. object) based on activity of vS1, tSTR and BLA cells during voluntary touch (0 to 5s). Up to 20 neurons were randomly chosen from each brain region for the SVM classifier. As a control we tested decoder accuracy when the context identity was shuffled in the 80% of object and social touch presentations (64 stimulations total) **i.** Decoder accuracy for touch context based on activity of 10 randomly selected cells in vS1, tSTR and BLA per animal across 50 ms bins throughout voluntary touch (−1 to +2 s).

We recorded the activity of single units across these three regions as mice were presented with 40 bouts of social and object touch under voluntary or forced conditions. Some neurons increased their firing in response to different presentations of touch, whereas others suppressed their firing (**Supplementary Fig. 4d**). We first considered the mean activity of all neurons, regardless of whether they were excited or suppressed by touch (**Fig. 3b**). On average, neurons across all three regions showed increased firing over baseline in response to both social and object touch, and this was apparent even before the platform stopped, because mice could initiate contact with their whiskers as the platform approached (**Fig. 3c**). Neurons in vS1 showed greater firing to social than object touch, whereas tSTR neurons showed stronger modulation to object touch (especially at the onset of touch) and BLA neurons did not show an obvious preference. Whether or not the animal could choose to engage in social touch also influenced neural activity in a region-dependent manner, as forced touch trials elicited a higher average response in both vS1 and tSTR than voluntary touch trials during the initial contact with the object (**Fig. 3d**; vS1 p=0.039, tSTR p=0.055).

Across the population of neurons in each region (e.g., vS1 in **Fig. 3b**), we observed a variety of response profiles: some neurons were suppressed by touch, some were excited transiently upon contact, others exhibited sustained firing during touch, and still others were barely modulated by touch. To compare the activity of units with such different behaviors, we performed principal component analysis (PCA) on trial-averaged activity followed by k-means clustering of the top PCA components (see Methods). This identified 5 significantly distinct clusters of single units in vS1, 6 clusters in tSTR and 4 clusters in BLA (**Fig. 3e**). Neurons that were least modulated by social/object touch (Cl. 1, Cl. 6, Cl. 12) tended to be the most abundant in all regions (**Supplementary Fig. 5a**). Surprisingly, neurons that were suppressed by social/object touch (Cl. 1-2, 6-7, 12-13) tended to represent a substantial proportion of the entire population (e.g., 47%, 36% and 66% in vS1, tSTR, BLA, respectively, for voluntary touch; **Supplementary Fig. 5)**.

Based on similarities in neural responses, we combined clusters that were strongly excited (Cl. 3-5 in vS1, Cl. 9-11 in tSTR, and Cl. 14 & 15 in BLA) or strongly suppressed (Cl. 2 in vS1, Cl. 7 in tSTR, and Cl. 13 in BLA) by touch within each region and focused on these for subsequent analyses. First, we compared differences in the modulation of their firing by touch context as an average of all 40 presentations (**Fig. 3f**). In vS1, we found that both excited and suppressed cells were preferentially modulated by social touch (**Fig. 3f**; **Supplementary Fig. 6**; p-value range: 0.07 to <0.001). In tSTR, excited cells also showed greater modulation by social touch for voluntary presentations (**Fig. 3f**; p=0.004), but the opposite occurred during forced interactions (**Fig. 3f**; **Supplementary Fig. 6b**; p=0.012). In the BLA, only excited neurons in Cl. 14 showed significantly higher modulation by forced object touch (**Supplementary Fig. 6b**; p<0.001). However, suppressed neurons in tSTR and BLA were not differentially modulated by social vs. object touch (**Supplementary Fig. 6**).

When we examined modulation by touch choice (voluntary vs. forced), we found that tSTR excited neurons showed greater modulation for forced object touch, when the object was in the animal’s personal space, which was the most aversive condition. Therefore, we also examined whether neural responses to forced object touch might differ during epochs when WT mice display aversion. We found that excited units in the tSTR (but not in other brain regions) were more active during bouts of avoidance/aversion compared to times when they did not display such behaviors (**Fig. 3g**; p=0.004).

These results show that: 1. vS1 neurons can discriminate touch context and are preferentially modulated by social touch; 2. tSTR neurons care about touch choice, because they are modulated in opposite ways by social vs. object touch depending on whether mice can choose to engage; 3. tSTR and BLA neurons are preferentially modulated by forced object touch, which produces more AFEs in WT mice. Thus, for vS1 and tSTR, being able to discriminate between social vs. object touch matters for the behavioral response to stimuli that require self-initiated exploration; whereas higher tSTR and BLA firing for a stimulus predicts avoidance/aversion when it is unwanted (forced).

### Touch context can be decoded from vS1, tSTR and BLA population activity

Considering the significant differences in neural activity between social vs. object touch, we hypothesized that SVM linear classifiers trained on Neuropixels data for all cells (see Methods) would accurately decode touch context. Indeed, we found high decoding accuracy for both voluntary and forced conditions, even with a handful of neurons (**Fig. 3h; Extended Data. Fig. 7a**). When estimating decoding accuracy across time, performance was always >60% and increased sharply for tSTR and vS1 upon contact (**Fig. 3i; Supplementary Fig. 7b**). Decoding performance was highest for those clusters that are most strongly excited by touch, such as Cl. 5 in vS1 and Cl. 10 in tSTR (**Supplementary Fig. 7c-e**).

### Forced touch recruits the highest proportion of object-preferring neurons in the BLA

The fact that neural activity in vS1, tSTR, and BLA showed differential modulation by social vs. object touch raises the possibility that certain neurons may exhibit a true preference to either object or social touch. We used receiver operating characteristic (ROC) analysis to categorize object-preferring and social-preferring cells (**Fig. 4a-c** and **Supplementary Fig. 8a-b**; see Methods). We found that at least 10% of units that were excited by touch showed a significant preference for social touch (greater than expected by chance; see Methods), irrespective of the brain region (**Fig. 4d**). In vS1, there was a similar proportion of object- and social-preferring units regardless of whether interactions were voluntary or forced (17-23%; **Fig. 4d-e**). In contrast, we found a higher proportion of object-preferring cells during forced interactions in the tSTR and in the BLA (**Fig. 4e**; p=0.055 and p=0.003, respectively). Together with the greater modulation of excited cells in tSTR and BLA by forced object touch, which trigger the most aversion in test mice, these findings further support the notion that these two regions are implicated in aversion to touch within the animal’s personal space (when it has no choice but to engage).

**Fig. 4:**
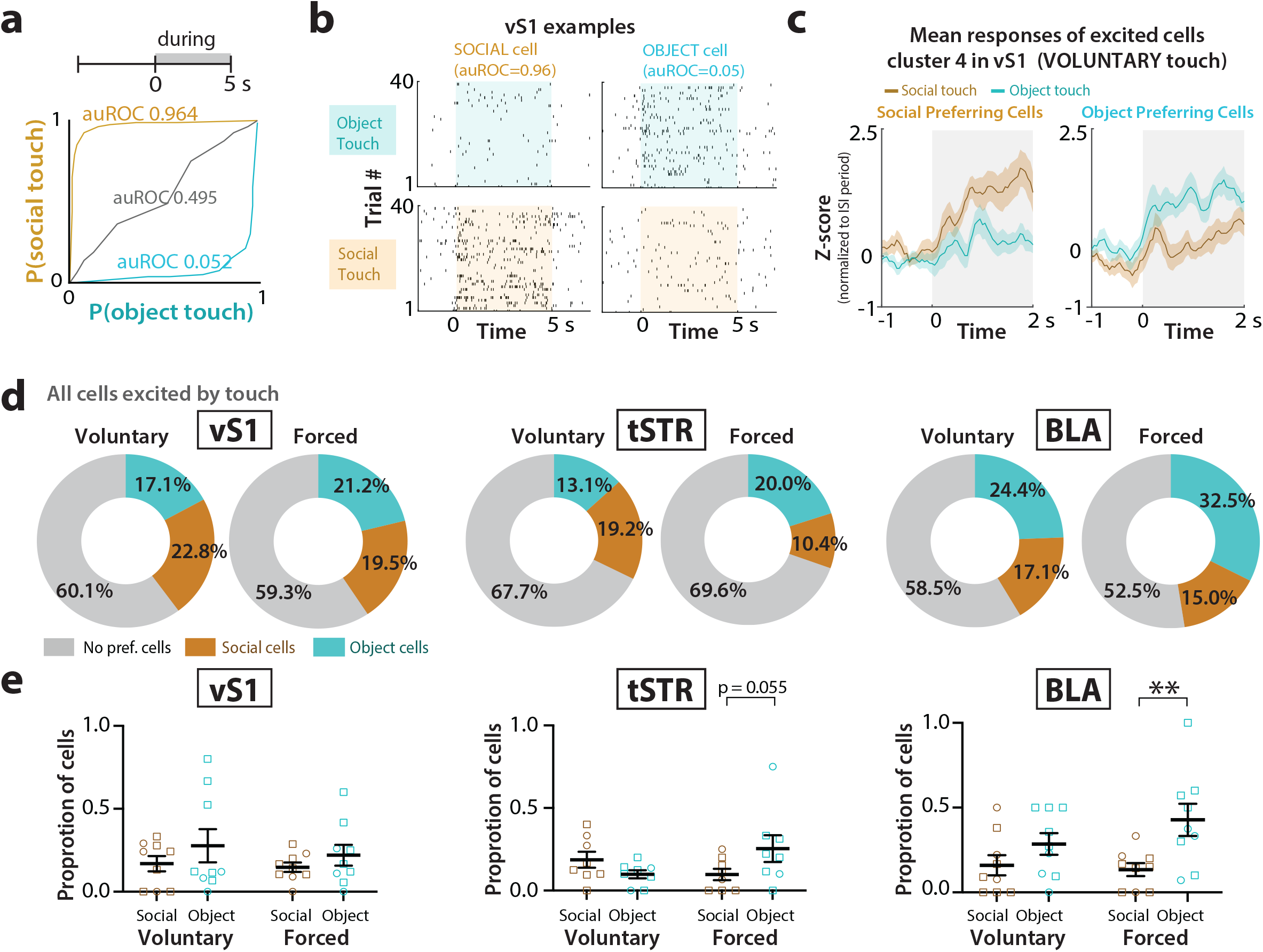
Higher proportion of object-preferring cells in tSTR and BLA. **a.** Example ROC curves (and corresponding ROC values) for a significant social-preferring cell, an object-preferring cell, and a neutral cell in vS1 (from cluster 4). **b.** Spike rasters across each presentation of social touch and object touch for the same example social-preferring and object-preferring cells in vS1 shown in panel *a*. **c.** Mean z-score firing rates of the same example social-preferring and object-preferring cells in vS1 during voluntary object touch and social touch. **d.** Percentage of object-preferring and social-preferring cells in vS1, tSTR and BLA during voluntary and forced touch as total of all cells. **e.** Proportion of object-preferring and social-preferring cells in vS1, tSTR and BLA for individual mice (n=9 WT mice). **p<0.01 for two-way ANOVA with Bonferroni’s. Squares=males, circles=females. 1 mouse was excluded according to ROUT’s analysis for tSTR.

### *Fmr1* KO mice do not discriminate social valence and find social interactions in their personal space more aversive than WT controls

To further understand the relationship between neural activity and behavioral responses to social vs. object touch, we next turned to *Fmr1* KO mice, because we recently discovered that they show similar aversion to social and object touch^21^. Thus, we hypothesized that neural activity may not be differentially modulated by touch context in this model of autism. We once again found that *Fmr1* KO mice (n=10 mice) manifest similar levels of avoidance running and AFEs to social and object touch (**Fig. 5a,b**; p>0.05 for all). When assessing how mice responded depending on whether they could choose to engage or not, we found that *Fmr1* KO mice displayed more aversive whisker protraction to object touch during forced presentations compared to voluntary interactions (**Fig. 5b**; p=0.009; **Supplementary Fig. 9a**; p=0.012).

**Fig. 5:**
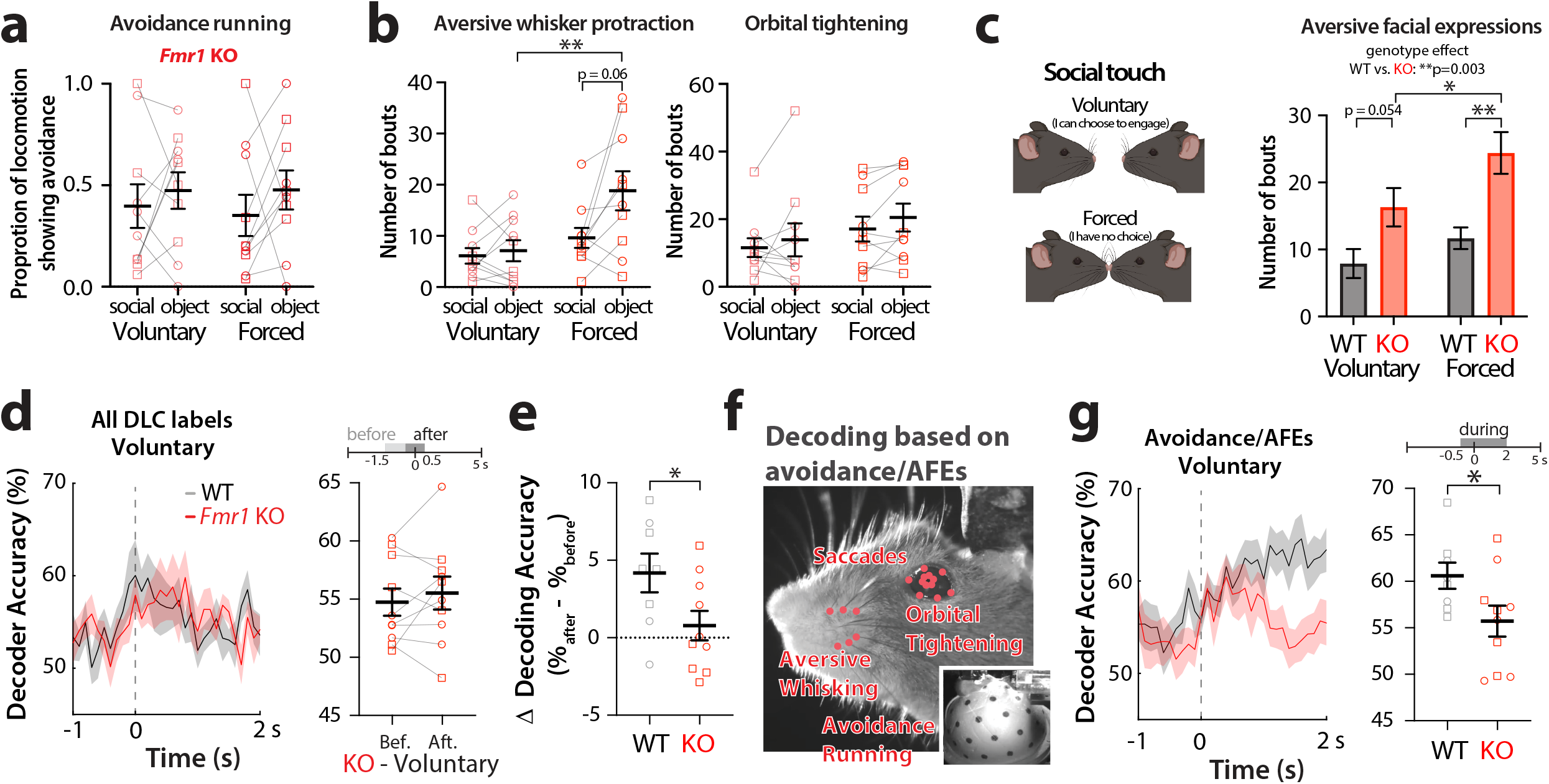
*Fmr1* KO mice show similar aversion to social and object touch and find social interactions within their personal space particularly aversive. **a.** Running avoidance during the first 5 presentations of touch in *Fmr1* KO mice (n=10) for social touch and object touch. **b.** Number of bouts of prolonged whisker protraction (left) and orbital tightening (right) in *Fmr1* KO mice across all 40 presentations of social or object touch in *Fmr1* KO mice. Squares=males, circles=females. **p<0.01 for two-way ANOVA with Bonferroni’s. **c.** Left: Cartoon of how test mice might perceive touch choice (voluntary vs. forced touch). Right: number of bouts of AFEs (aversive whisker protraction and orbital tightening) for *Fmr1* KO and WT mice during voluntary vs. forced social touch. **p<0.01 and *p<0.05 for two-way ANOVA with Bonferroni’s. **d.** Left: Decoder performance for touch context (social vs. object) using all DLC labels on the mouse’s face in *Fmr1* KO and WT mice across time (−1 to +2 s after platform stops). Right: Decoder accuracy for touch context before (−1.5 to −0.5 s) and after (−0.5 to +0.5 s) in *Fmr1KO* mice. Squares=males, circles=females. *p<0.01 for unpaired parametric t-test. **e.** Change in decoder accuracy (% after minus % before) for touch context using all DLC whisker labels in *Fmr1* KO and WT. Squares=males, circles=females. *p<0.05 for unpaired parametric t-test. **f.** Avoidance running, saccades, and AFEs were used to decode touch context with SVM classifiers. **g.** Left: Decoder accuracy for touch context based on aversive behaviors in WT and *Fmr1* KO mice (from −1 s to +2 s after platform stops). Right: Mean decoder accuracy in *Fmr1* KO than WT mice (−0.5 to +2 s) for individual mice. Squares=males, circles=females. *p<0.01 for unpaired parametric t-test.

When comparing across genotypes, we found *Fmr1* KO mice showed significantly more AFEs (whisker protraction & orbital tightening) to social touch than WT controls (**Fig. 5c**; p=0.003). Interestingly, AFEs were significantly greater in *Fmr1* KO mice for forced social touch than for voluntary social touch (**Fig. 5c**; p=0.003). Thus, social touch seems especially bothersome to *Fmr1* KO mice when it takes place within their personal space.

### Orofacial movements from *Fmr1* KO mice are not as good at decoding touch context as those of WT mice

Considering how *Fmr1* KO mice show similar behavioral responses to social and object touch (**Fig. 5a-b**), we hypothesized that SVM classifiers trained on their facial expressions would show lower accuracy in discriminating touch context compared to WT mice. Decoder performance from all DLC labels did not change in *Fmr1* KO mice following the movement of the platform towards the test mouse, unlike what we had observed for WT mice (**Fig. 5d-e**; KO p>0.05, WT vs *Fmr1* KO p=0.043). We wondered whether classifiers trained on aversive behaviors might also perform differently in both genotypes. We combined behavior data for running avoidance, AFEs, and eye saccades (associated with avoidance in ASD^48,49^), and the resulting decoder showed better performance for WT mice (**Fig. 5f-g**; p=0.046).

We also tested the performance of classifiers trained on orofacial movements from *Fmr1* KO mice at discriminating touch choice (voluntary vs. forced). Just as for WT mice, we observed a significant increase in decoder accuracy upon movement of the platform towards *Fmr1* KO mice (**Supplementary Fig. 9b-c**; p=0.032).

### vS1, tSTR and BLA neurons in *Fmr1* KO mice do not distinguish between voluntary social vs. object touch

We next used Neuropixels to record from units in vS1, tSTR and BLA of *Fmr1* KO mice, and used the same clustering approach. The proportion of units in various clusters differed between WT and *Fmr1* KO mice (**Supplementary Fig. 5**). For example, in vS1 there were significantly fewer cells in Cl. 2 and more in Cl. 5 in *Fmr1* KO mice, which are the clusters that, in WT mice, are the most modulated by social touch (**Supplementary Fig. 5b**). With regards to touch context, neurons in *Fmr1* KO mice responded similarly to social and non-social touch under voluntary conditions (**Fig. 6a, top**; **Supplementary Fig. 10a-b**). Indeed, when comparing the difference in modulation *(Δ modulation*) of excited neurons between social and object touch, we found that *Fmr1* KO mice uniformly had significantly smaller magnitudes across brain regions than WT mice (**Fig. 6a, bottom**; voluntary: vS1 excited p=0.013, tSTR excited p=0.038), which matches the behavioral responses under voluntary conditions. However, under forced touch conditions, neurons excited by touch in the tSTR and BLA of *Fmr1* KO mice were more strongly modulated by object touch than by social touch (**Fig. 6a**; **Supplementary Fig. 10b**), just as we had seen for WT mice (**Fig. 3e**).

**Fig. 6:**
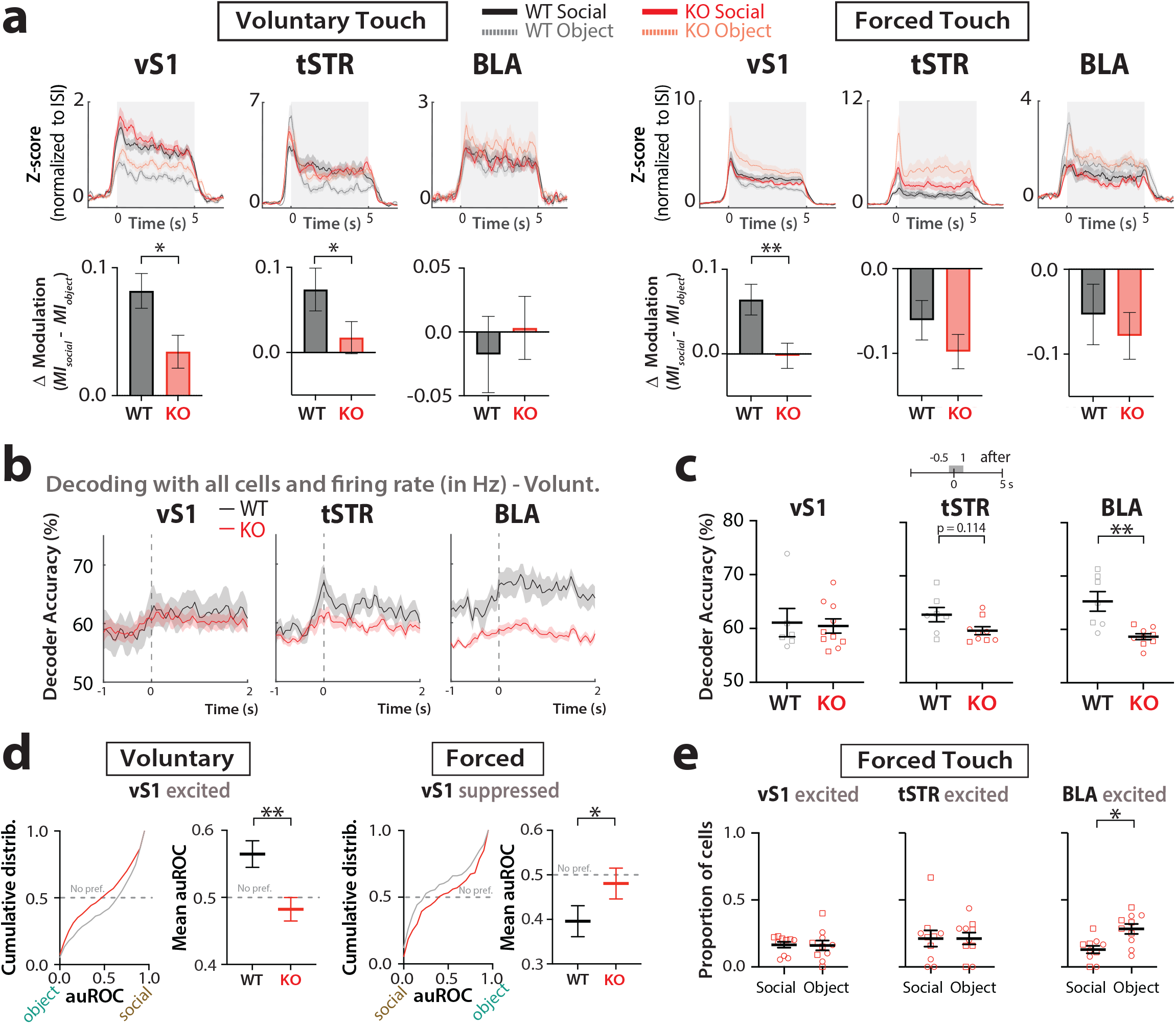
Lack of modulation of neural activity by touch context in *Fmr1* KO mice during voluntary presentations and reduced social preference compared to WT mice. **a.** Top: Z-score activity (normalized to ISI period) of excited cells in vS1, tSTR and BLA during social touch and object touch, for voluntary (left) and forced presentations (right). Bottom: Corresponding Δ modulation (*MI_social_* – *MI_object_*) by social vs. object touch of cells in vS1, tSTR and BLA of WT and *Fmr1* KO mice *p<0.05, **p<0.01 for unpaired nonparametric or parametric t-test. **b.** Decoder accuracy for touch context, averaged across mice, based on activity of 10 randomly selected cells in vS1, tSTR and BLA (50 ms time bins) during the presentation period of voluntary touch (−1 to +2 s). **c.** Mean decoder accuracy for touch context based on activity of 10 randomly selected cells in vS1, tSTR and BLA (−0.5 s to +1 s after that platform stops) for individual WT and *Fmr1* KO mice. Squares=males, circles=females. **p<0.01 for unpaired parametric t-test. **d.** Cumulative distribution (left) and mean auROC values (right) for excited and suppressed cells in vS1 (voluntary and forced touch, respectively) in WT and *Fmr1* KO mice. For excited cells auROC values above 0.5 correspond to social preference, while values below 0.5 correspond to an object preference; the opposite is true for suppressed cells; auROC values of 0.5 correspond to no preference. **p<0.01, *p<0.05 for unpaired parametric t-test. **e.** Proportion of object-preferring and social-preferring excited cells in vS1, tSTR and cells during forced touch in *Fmr1* KO mice. Squares=males, circles=females. *p<0.05 for paired parametric t-test.

In line with these results, SVM classifiers trained on neural data from the tSTR and BLA in *Fmr1* KO mice showed reduced decoding accuracy of touch context (under voluntary conditions) compared to WT mice (**Fig. 6b-c**; tSTR: p=0.114; BLA: p=0.001).

Overall, these results mirror our behavioral observations and argue that circuits from *Fmr1* KO mice fail to discriminate between social and object touch under voluntary conditions; however, when choice is removed (forced touch), *Fmr1* KO mice demonstrate greater modulation of neurons in BLA and tSTR by object touch, which is the most aversive.

### Neurons in vS1 of WT mice, but not *Fmr1* KO mice, show a preference for social touch

We next used ROC analyses to calculate the proportion of social-preferring and object-preferring excited neurons in vS1, tSTR and the BLA of *Fmr1* KO mice. When comparing the cumulative distributions of ROC values (auROC) for all excited neurons in vS1 we found a significantly lower social preference in *Fmr1* KO mice compared in WT controls (**Fig. 6d**; voluntary p=0.002, forced p=0.045). Only the BLA showed a significantly higher proportion of object-preferring cells (**Fig. 6e**; p=0.010), just as we had seen in WT mice (**Fig. 4d**). Thus, vS1 of *Fmr1* KO mice seems generally unable to differentiate between social and non-social stimuli.

## DISCUSSION

The main goal of this study was to investigate how different brain circuits are modulated by social facial touch and how this relates to aversive responses. We focused on both the context of touch (the social valence of touch) and the importance of having a choice to engage in social touch (the notion of peri-personal space).

Our main findings can be summarized as follows (**Supplementary Fig. 11**): 1. WT mice tolerate social touch but show aversion to object touch, especially at close range (forced touch). 2. vS1 neurons care about touch context and are preferentially modulated by social touch. 3. In contrast, tSTR and BLA neurons care most about touch choice, firing preferentially to object touch when it occurs in the animal’s personal space. Because forced object touch elicited more avoidance/AFEs, we surmise that activity in BLA and tSTR relates to behavioral responses when the animal is forced to interact; 4. *Fmr1* KO mice show similar degrees of avoidance/AFEs to both social and object touch, which means they are unable to distinguish, at the behavioral level, the social valence of an interaction; 5. Consistent with this, neural activity in all three regions in *Fmr1* KO mice was less able to discriminate between social vs. object touch. Moreover, *Fmr1* KO mice showed striking aversion to forced touch (particularly social touch), and activity in the BLA tracked this, suggesting that maladaptive responses to stimuli in the animal’s peri-personal space map onto amygdalar circuits.

Our cFos expression data suggests that processing of social touch involves circuits distributed across many brain regions. We have identified several interesting brain regions worth exploring in the context of social touch. Notably, regions such as the ICx, PAG, or the PVHy have already been implicated in social behaviors^50–53^. Future studies should assess whether these other brain regions can also distinguish social from non-social tactile inputs, or whether this is dependent on an animal ability’s to voluntarily engage in touch.

The fact vS1 neurons still showed a social preference when the animal has no choice but to engage, suggests that the valence of the social stimulus matters to cortical neurons. Thus, vS1 neurons are not solely responding to a difference in shape/texture between a plastic tube and a mouse, but also care about valence, presumably relying on inputs from other brain regions that encode pleasurable aspects of social touch^18–20^. Furthermore, the neural activity in vS1, but not the activity of tSTR and BLA, aligns well with how WT mice tolerate social touch. Without this neural discriminability, we surmise that animals would no longer differentiate social from non-social stimuli behaviorally, as occurs for *Fmr1* KO mice (see below).

Unlike vS1, activity in the BLA showed no preference for social or object touch during voluntary interactions. Thus, the social valence of a stimulus does not seem to matter to BLA neurons, at least in our assay. This was surprising considering the roles that have been ascribed to BLA in social exploration and approach^25,46^. Instead, BLA neurons showed a preference for the most aversive stimulus, forced object touch (and a greater percentage of object-preferring cells) in mice of both genotypes. We interpret this to mean that touch within personal space matters to the BLA, and that aversive stimuli are represented in amygdalar circuits^26,54^. The BLA does receive projections from the ACCx and has been linked to learning of aversive sensory stimuli^26,29^ and with the encoding of exploratory and aversive behavioral states^24^.

Activity in the tSTR was modulated in opposite ways by social vs. object touch depending on whether the interaction was forced or voluntary. It implies that neurons in the tSTR, unlike those in vS1, care less about stimulus context (social valence) and more about whether the interaction takes place within personal space (i.e., no choice but to interact) and about the ultimate behavioral response. Indeed, tSTR neurons were uniquely modulated during bouts of AFEs. Interestingly, tSTR receives inputs from amygdalar nuclei and prefrontal cortex^46,55^.

In contrast to WT mice, *Fmr1* KO mice display equal aversion to both social and object presentations, and therefore fail to recognize the social valence of a visitor mouse. This is consistent with the social disinterest that *Fmr1* KO mice show in a social interaction test^56^. Furthermore, we find that having a choice to engage with another mouse matters, and that *Fmr1* KO mice (but not WT mice) find forced social touch at close proximity more aversive compared to when they voluntarily initiate social touch from a distance. Thus, social interactions that invade personal space may be especially bothersome in autism. This is perhaps the first demonstration of behavioral responses related to peri-personal space, a topic of increasing interest in the autism field^54,57,58^.

Also unlike WT mice, cortical and striatal circuits of *Fmr1* KO mice were not preferentially modulated by either social or non-social stimuli during voluntary interactions (also reflected in the reduced decoding accuracy for touch context based on neural activity). We therefore conclude that a failure to discriminate social from non-social touch at the circuit level could explain the reduced social interest in autism. We previously demonstrated that the maternal immune activation model of autism shows similar aversion to social touch as *Fmr1* KO mice^21^. It will be important to investigate how cortical, striatal, and amygdalar circuits in this and other models respond to social touch. Moreover, chemogenetic or optogenetic manipulations within these regions would help establish causal links between these circuits and social avoidance, which could be important for the development of strategies that alleviate social deficits in autism.

## ONLINE METHODS

### Animals

Adult male and female C5BL/6 mice at postnatal day 60-90 were used for all experiments. A cohort of adult mice (9 male and 8 female) were used for TRAP labeling of neurons activated by social/object touch. These so-called ‘TRAP’ mice were obtained by crossing Fos^2A-iCreER/+^ (TRAP2) (JAX line 021882) with R26^Ai^^14/+^ (Ai14) (JAX line 030323).

A second cohort of mice (>20 g in weight) was used for electrophysiological recordings and were derived from the following mouse lines based on prior publications: wildtype (WT) B6J (JAX line 000664), *Fmr1* KO (JAX line 003025)^32,33,59–61^. In total 9 WT (6 male and 3 female), and *Fmr1* KO mice (6 male, and 4 female) were used for Neuropixels recordings.

All mice were group-housed with access to food and water using HydroGel (ClearH_2_O) *ad libitum* under a 12 hour light cycle (12 hours light/12 hours dark) in controlled temperature conditions. Mice with Neuropixels implants were single housed during habituation and behavioral testing to avoid damage to the implant by group housing with other animals (∼1-2 weeks). All experiments were done in the light cycle and followed the U.S. National Institutes of Health guidelines for animal research under an animal use protocol (ARC #2007-035) approved by the Chancellor’s Animal Research Committee and Office for Animal Research Oversight at the University of California, Los Angeles.

### TRAP2 mice drug preparation

4-hydroxytamoxifen (4-OHT, Sigma-Aldrich #H6278) was dissolved in 20 mg/ml in ethanol and was aliquoted and stored at −20°C up to 4 weeks. On the day before behavioral testing, 4-OHT was redissolved in 70% EtOH by warming the aliquot at 37°C and vortexing vigorously for 1 min 2-3 times. Corn oil (Sigma Aldrich, C8267) was added to each aliquot for a final concentration of 10 mg/ml of 4-OHT. The aliquots were vacuum centrifuged for 30 min until all EtOH was evaporated. 4-OHT was stored in 4°C for next-day use.

### Behavioral social/object touch experiments for TRAP labeling

To identify brain regions involved in social touch, we used the behavioral assay we recently developed ^21^. Adult TRAP2 mice (and the corresponding visitor mice in the social touch assay) were first surgically implanted with a titanium head bar. Briefly, mice were anesthetized with isoflurane (5% induction, 1.5-2% maintenance via nose cone v/v) and secured on a motorized stereotaxic frame (Kopf; StereoDrive, Neurostar) via metal ear bars. The head of the animal was shaved with an electric razor and the skin overlying the skull was then sterilized with three alternating swabs of 70% EtOH and betadine. A 1 cm long midline scalp incision was made with a scalpel and the custom U-shaped head bar (3.15 mm wide x 10 mm long) was secured on the back of the skull first with Krazy Glue and then with a thin layer of C&B Metabond (Parkell) applied to the dry skull surface. The entire skull was then covered with acrylic dental cement (Lang Dental). This surgery lasted ∼15-20 min and mice fully recovered within 30 min, after which they were returned to group-housed cages.

At least 48 h after head bar implantation, TRAP2 mice were habituated to head restraint, to running on an air-suspended 200 mm polystyrene ball, and to the movement of a motorized stage that was used for repeated presentations of an inanimate object or a stranger mouse. The stage consisted of an aluminum bread board (15 x 7.6 x 1 cm) attached to a translational motor (Zaber Technologies, X-LSM100A), the movement of which was fully controlled through MATLAB (Mathworks). All of this occurred in a custom-built, sound-attenuated behavioral rig (93 x 93 x 57 cm) that was dimly illuminated by two infrared lights (Bosch, 850 nm). For habituation of TRAP2 mice, test mice were placed on the ball for 20 min each day for 14 consecutive days before testing. In parallel, ‘visitor’ mice (stranger to the test mouse) were habituated to head-restraint in a plexiglass tube (diameter: 4 cm) on the motorized stage. The stage translated at a constant speed of 1.65 cm/s. In its neutral starting position, the snout of the visitor mouse was 6 cm away from that of the test mouse.

Following habituation, all TRAP2 test mice were single-housed the day before the social touch assay^21^. On the day of behavioral testing and 30 min prior to testing, TRAP2 mice were injected with 4-OHT (50 mg/kg, i.p.). TRAP2 test mice were tested under three different conditions: 1. no touch, in which the platform moved back and forth in repeated bouts but was empty; 2. object touch, in which test mice experienced repeated bouts of forced interactions with a plastic 50 mL Falcon conical tube; and 3. social touch, in which test mice experienced repeated bouts of forced interactions with a visitor novel mouse (stranger to the test mouse). For the forced object/social interactions, the stage stopped at a position that brought the tip of the plastic tube or the snout of the visitor mouse in direct contact with the snout of the test mouse. These positions were calibrated before each experiment. For the condition where the platform was empty, the stage was moved to a set template position that was tested during calibration for forced interactions. Each bout lasted 5 s, with a 5 s interstimulus interval (ISI) during which the platform moved away by 1 cm and the object/mouse was out of reach of the test mouse’s whiskers. The ISI included a 1.2 s period of back-and-forth travel time for the platform. Each session (no touch, social touch, object touch) lasted 30 min, which was equivalent to a total of 180 such presentations.

Following the assay, TRAP2 test mice were returned to their single cage housing until the end of the day (∼6-8 h), at which point they were placed back in group housing, and then they were perfused at 72 h. For each session of behavioral testing, at least 3 mice were used (one mouse each for no touch, object touch and social touch condition).

### Histology and quantification of cFos expression in TRAP2 mice

72 h after 4-OHT induction (to allow *Cre* recombination to occur), TRAP2 mice were transcardially perfused with 4% paraformaldehyde (PF) in cold PBS (0.1M) and their brains were harvested and left overnight in 4% PF. Next, fixed brains were sliced coronally to obtain 60 μm sections. The coronal sections were mounted on VectaShield glass slides and stained with DAPI (Vector Laboratories). Sections were imaged on an Apotome2 microscope (Zeiss; 10x objective). Images were taken as a z-stack ranging from 30-50 μm (Zen2 software, Zeiss). ImageJ was used to quantify the density of cells expressing tdTom in each brain region (cFos-tdTomato^+^ cells/mm^2^). Cell densities in each brain region from ‘object touch’ mice (TRAP2-OBJECT) and ‘social touch’ mice (TRAP2-SOCIAL) were normalized to averaged cell density in each brain region from ‘no touch’ mice (TRAP2-CTRL) in the same session of behavioral testing to account for variability in tamoxifen preparation from one session to the next.

### Surgical implantation of Neuropixels probes for chronic recordings

Each Neuropixels 1.0 probe (Imec, PRB_1_4_0480_1_C) was first connected to the acquisition hardware to confirm that the probe was functional using the SpikeGLX data acquisition software (see below) both prior to and after soldering a grounding wire (0.01 inches, A.M. Systems) to the ground and reference pads in the probe flex cable. The probe was inserted and screwed into a dovetail probe holder (Imec, HOLDER_1000_C) and set aside for surgical implantation. A custom-made external chassis cover (eventually used to protect the probe during implantation) was 3D-printed (Hubs) using standard black resin (Formlabs, RS-F2-GPBK-04). The CAD files for the 3D printed cover were acquired at https://github.com/Brody-Lab/chronic_neuropixels^62^ - we used the external casing part.

Adult mice were anesthetized with isoflurane and placed on a motorized stereotaxic frame. Their head was shaved and the scalp sterilized as above. A 1 cm long midline scalp incision was made with a scalpel and a small burr hole (0.5-0.8 mm diameter) was drilled over the cerebellum (2 mm posterior to Lambda) with a dental drill (Midwest Tradition) through which a stainless-steel ground screw (M1, McMaster Carr) was loosely screwed. A second burr hole (0.5-1 mm diameter) was drilled at the probe implantation site at coordinates −1.46 AP, 2.9 ML 3.75 DV (in mm). This allowed for targeting of vS1, tSTR and BLA simultaneously with a single shank Neuropixels probe. Saline soaked Surgifoam (Ethicon) was placed in both craniotomies while a thin layer of C&B Metabond (Parkell) was applied to the dry skull surface. A small well (0.75 cm diameter, 1 cm height) was built around the craniotomy site with self-adhesive resin cement (RelyX Unicem 2 Automix, 3M ESPE) and set with dental curing lamp (Sino Dental). Surgifoam was removed from the implantation craniotomy and saline was applied to maintain tissue hydration.

Before insertion, the probe holder (with the probe attached) was screwed and secured to a micromanipulator (StereoDrive, Neurostar). To enable histological reconstruction of the probe tract, the tip of the shank was dipped for 30 s in DiI (in 1-2 mg/mL in isopropyl alcohol; Sigma Aldrich, applied onto Parafilm, Bemis). DiI fluorescence enabled subsequent histological reconstruction of the probe tract in fixed tissue sections. The ground wire soldered to the probe was then wrapped around the ground screw, as it was being screwed into the ground burr hole. Conductive epoxy (8331, MG Chemical) was applied on the ground screw and wire. A titanium U-shaped head bar (3.15 x 10 mm) was affixed to the skull with Metabond (Parkell) caudal to the ground screw, to allow for head-restraint during recordings and behavioral testing. Next, the probe shank was lowered at a rate of 10 μm/s with StereoDrive through the implantation craniotomy. Saline in the cement well was then absorbed carefully with Surgifoam and replaced with Dura-Gel (Cambridge Neurotech). Additional light curable resin cement was applied and cured to the cement well and onto the probe base (avoiding contact with the shank). An additional layer of Metabond was applied on the skull, including on the ground craniotomy site and along the outside of the resin cement well. The two parts of external chassis were then placed around the probe and cemented together and to the resin cement wall with acrylic (Lang Dental). This surgery lasts 3-4 h and hydration was provided by injecting saline every hour (0.1 mL, i.p.). Mice fully recovered within 1-2 h after surgery. Afterwards, implanted animals were single housed for habituation and behavioral testing. Mice were given carprofen (Rimadyl) immediately after surgery and again at 24 h and 48 h post-op, and given ad lib access to HydroGel (ClearH_2_O) and food.

### Social touch assay in mice with chronic Neuropixels implants

Following probe implantation, test mice were subjected to the social touch assay described above, but in addition to forced object/social touch we introduced additional interactions (see below). First, mice bearing Neuropixels implants were habituated to head restraint, to running on the polystyrene ball, and to the behavioral apparatus (just as for the TRAP experiments above, but for only 7-9 d).

Following habituation, test mice were subjected to both voluntary and forced interactions with a visitor mouse or a novel inanimate object over the course of 2 d. On day 1, test mice were placed on the ball and recorded for a 2 min baseline period (the plexiglass tube on the moving stage was empty). Next, we inserted the plastic object (50 mL Falcon conical tube) into the plexiglass tube on the motorized stage. For this control interaction, the test mouse first experienced a 2 min period of no touch but was able to visualize the object in the neutral position (before touch, 6 cm away). Next, the motorized stage moved the object to within whisker reach of the test mouse for a total of 40 such presentations of either voluntary (whisker-to-object) or forced (snout-to-object) object touch. Each bout lasted 5 s, with a 5 s ISI during which the platform moved away by 1 cm and the object was out of reach of the test mouse. The ISI included the total travel time of the platform, 1.2 s.

After this object touch session, the test mouse was returned to its cage to rest for at least 1 h before being head-restrained again to undergo either voluntary or forced social touch session (same type of touch as previous session for object touch) with a visitor mouse. A same-sex, same age (P60-90) novel WT mouse was head-restrained inside the plexiglass tube on the motorized stage. Following a 2 min period in the neutral position where the test mouse could see but not touch the stranger mouse, the motorized stage moved to the position for voluntary social touch (whisker-to-whisker) or forced social touch (snout-to-snout) for 40 bouts (also lasting 5 s with a 5 s ISI where the mouse on the platform moved out of reach of the test mouse). The test mouse was then returned to its cage for at least 24 h.

On day #2 of behavior testing, the mouse was placed back on the ball again for a 2 min baseline period followed by a 2 min period of no touch. Depending on what interaction the test mouse had received voluntary or forced object and social touch on testing day #1, the mouse received 40 presentations of the alternate touch context with a novel object and another stranger mouse.

### Electrophysiological recordings

During the social touch behavioral assay and tactile defensiveness assay, electrophysiological recordings were performed using Neuropixels 1.0 acquisition hardware (Imec). The acquisition hardware was used in combination with PCI eXtensions for Instrumentation (PXI) hardware (PXIe-1071 chassis, PXIe-8381 remote control module and PXIe-6341 I/O module for recording analog and digital inputs, National Instruments). SpikeGLX software was used to acquire data (https://github.com/billkarsh/SpikeGLX, HHMI/Janelia Research Campus). Recording channels acquired electrical signals from the most dorsal region of the vibrissal primary somatosensory cortex (vS1) down to the most ventral region of the basolateal amygdala (BLA) using the deepest of the 960 electrode sites. Electrophysiological signals were processed with Kilosort2.5 (https://github.com/MouseLand/Kilosort) using default parameters for spike sorting and then manually curated with Phy2 (https://github.com/cortex-lab/phy)^63^. Only well isolated single units were used for electrophysiological data analysis. After manual curation, we selected units that passed the following criteria: ISI violation <10%, amplitude cutoff <10% and median amplitude >50 μV, as previously described^64^.

### Removal of Neuropixels probes

Neuropixels probes were explanted for subsequent re-use. Mice implanted with Neuropixels were anesthetized with isoflurane and secured on a stereotaxic frame. The external chassis was removed with a dental drill, and any excess acrylic or resin cement around the probe dovetail was gently drilled off while avoiding direct contact with the probe. The dovetail holder was inserted and screwed into the probe and attached to the stereotaxic arm. Resin cement was carefully drilled around the circumference of the resin cement well to separate the skull from the probe. Once the skull and probe were separated, the probe was lifted up using the motorized micromanipulator, until the probe was completely outside the resin cement well. The probe was removed from the dovetail holder and forceps were used to gently remove any excess resin from the probe. After explantation, the probe shank was fully immersed in 1% tergazyme (Alconox) for 24-48 h, followed by a 1-2 h rinse in distilled water.

### Histology and fluorescence imaging of probe location

Following probe removal, mice were anesthetized with 5% isoflurane and transcardially perfused with 4% PF and post-fixed overnight. At least 24 h after perfusion, the fixed brain was rinsed with PBS and sliced coronally with a vibratome to generate 50 μm sections. The coronal sections are mounted on slides with VectaShield mounting medium (Vector Laboratories). DiI fluorescence in each section per brain was imaged on an Apotome2 microscope (Zen2 software, Zeiss; 5x objective, 5×5 grid of images). ImageJ was used to visualize each section image and reconstruct the entry point of the probe shank to the tip of the probe shank in the brain.

### Electrophysiological data analysis

We first converted action potential spikes to firing rate estimates (in spikes per second) for each single unit by binning spike counts in 50 ms bins and dividing by the bin size. To generate peristimulus time histograms (PSTH), we smoothed firing rates with a 250 ms moving window and then averaged all touch presentations from 2 s before the onset of touch to 2 s after the end of touch [-2 to +7s]. Units were assigned as belonging to vS1, tSTR or BLA based on the dynamics of their action potential spiking by depth and across time (**Supplementary Fig. 4**). Regular spiking (RS) units in vS1 were identified based on the duration of their spike waveform (≥ 400 µs peak to trough).

### Classification of single unit responses to social and object touch

Despite the heterogeneity of single unit responses to voluntary and forced touch, we sought to determine whether some units behaved similarly to other, i.e., whether there exist different functional groups of neurons in each brain region. We performed clustering of single units twice using PSTHs of all presentations (object and social) of (1) voluntary touch and (2) forced touch. Clustering of the PSTHs was also done separately for each brain region (vS1, tSTR and BLA), so the procedure was employed 6 times. The clustering procedure we used takes the z-scored, trial-averaged PSTH of each unit and combines all responses into a matrix (PSTH x unit). Units from wild type and *Fmr1* knockout (KO) mice were included together within the PSTH x unit matrix. Principal component analysis (PCA) was then performed on this matrix followed by k-means clustering of the top *k* components that explained >95% of the variance. The gap statistic criterion was used to estimate the ideal number of clusters for each clustering followed by visual inspection of temporal firing of units in each cluster (to confirm their different responses). We applied 1,000 iterations of k-means clustering for each clustering procedure performed. Clustering of single units was also performed on vS1, tSTR and BLA units separately for the recording session in which the test mouse received forced touch from an inanimate toy mouse and similarly for the session in which the test mouse received forced touch from an anaesthetized mouse. The PSTHs were split by brain region (vS1, tSTR or BLA) and grouped as either voluntary (average of all trials of voluntary object and social touch) or forced (average of all trials of forced object and social touch). By grouping units in this manner, we could compare how a unit assigned to a cluster by k-means responds differentially to voluntary object and social touch and responds differentially to forced object and social touch.

### Modulation of single units by social and object touch

To quantify differences in the mean firing rates of units in each cluster between voluntary or forced object and social touch, we calculated the z-score firing rate normalized to the average firing rate during the ISI period. To assess the modulation of units by social and object touch we grouped together neurons from clusters with similar temporal properties. Units in clusters that were moderately to strongly excited were grouped together (‘excited’ cells), as were moderately or strongly suppressed units (‘suppressed’ cells). A modulation index (*MI*) was used to calculate how much the firing rate (*FR*) of each unit changed during the stimulus period (*stim*) relative to the ISI period in each trial:

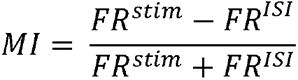

The MI was calculated using three different time ranges for the stim and ISI period. For calculating the MI over the entire stimulation period (*MI^STIM^*) we used the mean FR over 5 s during which the platform was stopped (touch) for *FR^stim^* and the mean firing rate over the 5 s ISI for *FR^ISI^*. For the MI of the first few seconds of the presentation period (*MI^SHORTSTIM^*), we used the mean FR from the first 3 s of presentation for *FR^stim^* and the mean FR for the 3 s prior to the presentation onset for *FR^ISI^*. For the MI during the period that the platform moves (*MI^PLATFORM^*), with the platform as the stim, we used the mean FR from [-1 1] s with time 0 as the presentation onset *FR^stim^* and the mean FR from [−3 −1] s for *FR^ISI^*. *MI^STIM^* was used to compare modulation of vS1 suppressed and excited cells, tSTR excited cells and BLA excited cells. We also assessed MI of units in each cluster. For assessing modulation by cluster, *MI^PLATFORM^* was used for units in clusters that showed a change in FR during the initial onset of touch, *MI^STIM^* was used for units in clusters that showed a sustained change in FR during the period of touch and *MI^SHORTSTIM^* was used for units in clusters that showed a larger change in FR the initial onset of touch as well as a sustained change in FR during touch.

### Single neuron coding of stimulus preference and behavior

To determine which units show clear preference for object vs. social touch, we used receiver operating characteristic (ROC) analysis, which was applied to the firing rate (Hz) during the presentation period [0, 5s], as previously described ^65–67^. Each unit’s preference was calculated based on the firing rate response to each trial relative to the mean PSTHs for object touch and social touch trials. Each trial was assigned a decision variable (DV) score and the DV for social touch and object touch trials was calculated as follows:

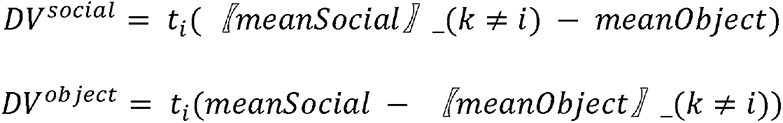

where *t_i_* is the firing rate for the current (*i*^th^ trial) and *meanSocial* and *meanObject* correspond to the mean social and object touch PSTHs. An ROC curve was obtained by varying the criterion value for the DV and the area under ROC (auROC) was calculated from the ROC curve using the MATLAB function *trapz.* The auROC value was considered significant by bootstrapping 1,000 times with a threshold probability of 0.05. Single units that were excited by touch and showed significant auROC values >0.5 were deemed to show preference for social touch (social cells) and those with significant values <0.5 showed preference for object touch (object cells). For suppressed units, those with significant auROC values <0.5 were social cells, and those with significant values >0.5 show were object cells. Units with non-significant auROC values were considered as showing no-preference.

### Decoding touch context from neuronal activity and behavior

We used support vector machine (SVM) linear classifiers to determine how well all neurons, or neurons within a particular cluster, could decode the presentation type (object vs. social) under voluntary or forced conditions. We used activity from 80% of trials (64/80 of both object and social touch trials) as the training dataset and the remaining 20% (26/80) was used for testing the classifier’s accuracy. Firing rates, either as an average of the stimulus period (0 to +5 s) or binned as 50 ms across the stimulus period (−2 to +7 s), were used as the feature space of the SVM. 100 iterations of the decoding analysis were performed in which the neurons of a cluster and trials were randomly chosen for the training and test data set. The mean decoding accuracy was calculated based on the average performance of all 100 iterations. In addition, we separately trained the decoder on neural data in which the trial labels were randomly shuffled for the test dataset (“Shuffled”). Decoding was performed with neurons within each cluster by brain region. A different population size of neurons ranging from 1 neuron to the 20 neurons was used for decoding context from averaged activity during the stimulus period and 20 neurons were used for decoding context across time. For decoding context (object vs social) or choice (voluntary vs forced) from behavior across time, we binned each behavioral measure in 100 ms bins. For decoding context from facial motion, a total 23 DeepLabCut labels were used and running avoidance, eye area, saccade direction and whisker protraction for decoding context from avoidance and aversive behaviors (see analysis of behavior below).

### Data analysis of behavioral data

During the social touch behavioral assay and tactile defensiveness assay, high-resolution videos (.avi files) were recorded of the test mouse’s eye, face, and body using three cameras (Teledyne Flir, Blackfly S USB3) at 120 FPS for behavioral analyses. Locomotion and running direction, facial expressions (including aversive facial expressions; AFEs) and pupil saccades were analyzed from these videos of the eye, face, and body. Running direction and AFEs (orbital tightening and whisker protraction) were quantified as described previously^21^ using MATLAB, FaceMap and DeepLabCut^21,35,68^. For locomotion, FaceMap was used to track the motion energy of polystyrene ball as the animal was running^35^. For analysis of pupil saccades and facial motion, a DeepLabCut neural network was trained on images from the eye or face videos to identify markers on the mouse’s pupil, eye, 6 whisker follicles, mouth and nose. The markers on the animal’s eye or face were used to quantify the following metrics: motion energy of each marker (change in marker position every 2 frames), saccades along temporal-nasal plane (displacement of pupil on x-axis), and eye area (pixel area of eye markers)^69^. For analysis of pupil saccades and facial motion and expressions (including AFEs), we excluded video frames when the animal was blinking, grooming, or other movements obscured the animal’s face.

### Statistical analyses

Statistical tests were performed in Prism software (GraphPad). Statistical analyses of normality (Lilliefors and Shapiro Wilk tests) were performed on each data set; if data deviated from normality (p<0.05) or not (p>0.05), appropriate non-parametric and parametric tests were performed. For parametric two-group comparisons, a Student’s t-test (paired or unpaired) was used. For non-parametric tests, we used Mann-Whitney test (two groups) and the Kruskal-Wallis test (repeated measures). Multiple comparisons across touch conditions and genotypes/groups were analyzed using two-way ANOVA with post-hoc Bonferroni’s test. If data was non-normal, we applied a logarithmic transformation on the data and compared the two-way ANOVA with and without the transformation. Since the statistical output of the two-way ANOVA was similar for the transformed and the non-transformed (non-normal data), we used the latter. All experiments were conducted in animals from at least two different litters for each genotype/group. For the figures related to the TRAP experiments, we used the number of tissue sections (with the same number per animal) as the sample size. For the remaining figures, the statistics were done on either the number units as the sample size or using individual mice as the sample size (averaged over cells for different mice) superimposed on individual data points. Because there are important sex differences in both the prevalence and symptoms of ASD^70,71^, we distinguished males from females across all figures (squares depict males, circles depict females). In all figures, the error bars denote standard error of mean (s.e.m.). Robust regression and outlier removal (ROUT) analysis was used to exclude outliers for data represented as individual mice. One WT animal was excluded from behavioral decoding as videos were not synchronized across cameras.

## Data and code availability

Custom code written in MATLAB for analysis of electrophysiological neural data and behavior is available at https://github.com/porteralab.

## ACKNOWLEDGEMENTS

We are grateful to Weizhe Hong, Lukas Oesch, and Anne Churchland for advice on analysis of behavioral and neural data. We thank Will Zeiger and Sotiris Masmanidis for comments on the manuscript. This work was supported by the following grants: R01NS117597 (NIH-NINDS), R01HD108370 and R01HD054453 (NIH-NICHD), Department of Defense (DOD, 13196175) awarded to C.P-C, Training in Neurotechnology Translation T32NS115753 (NIH), F31HD108043 (NIH/NICHD), and a graduate student fellowship from the Achievement Rewards for College Scientists Foundation to T.C., and the CARE Fellows Program to A.H. Some figure panel cartoons were generated with BioRender.

## SUPPLEMENTARY FIGURE LEGENDS

**Supplementary Fig. 1:**
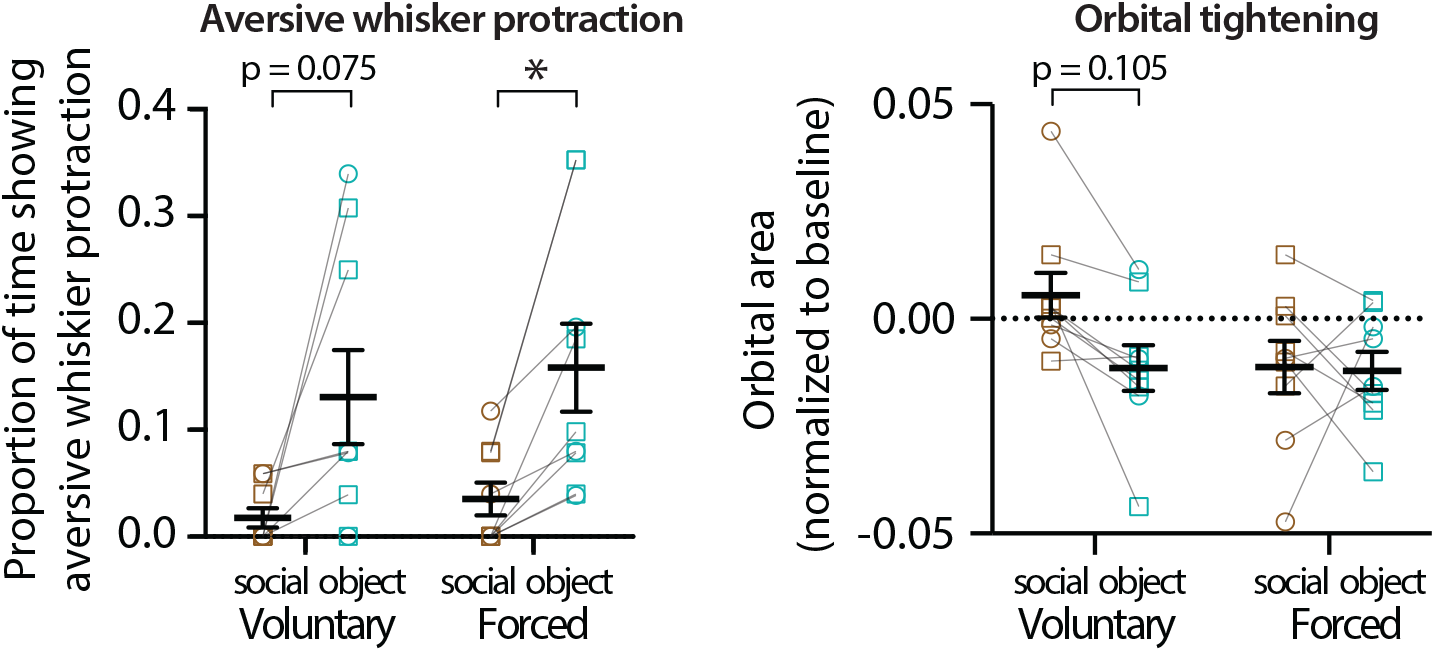
WT mice display aversive whisker protraction to forced object touch but not social touch. Proportion of time that WT mice manifest sustained aversive whisker protraction (left) and orbital area (right) in response social touch (brown) or object touch (cyan), over the first 5 voluntary or forced interactions. Squares=males, circles females. *p<0.05, two-way ANOVA with Bonferroni’s.

**Supplementary Fig. 2:**
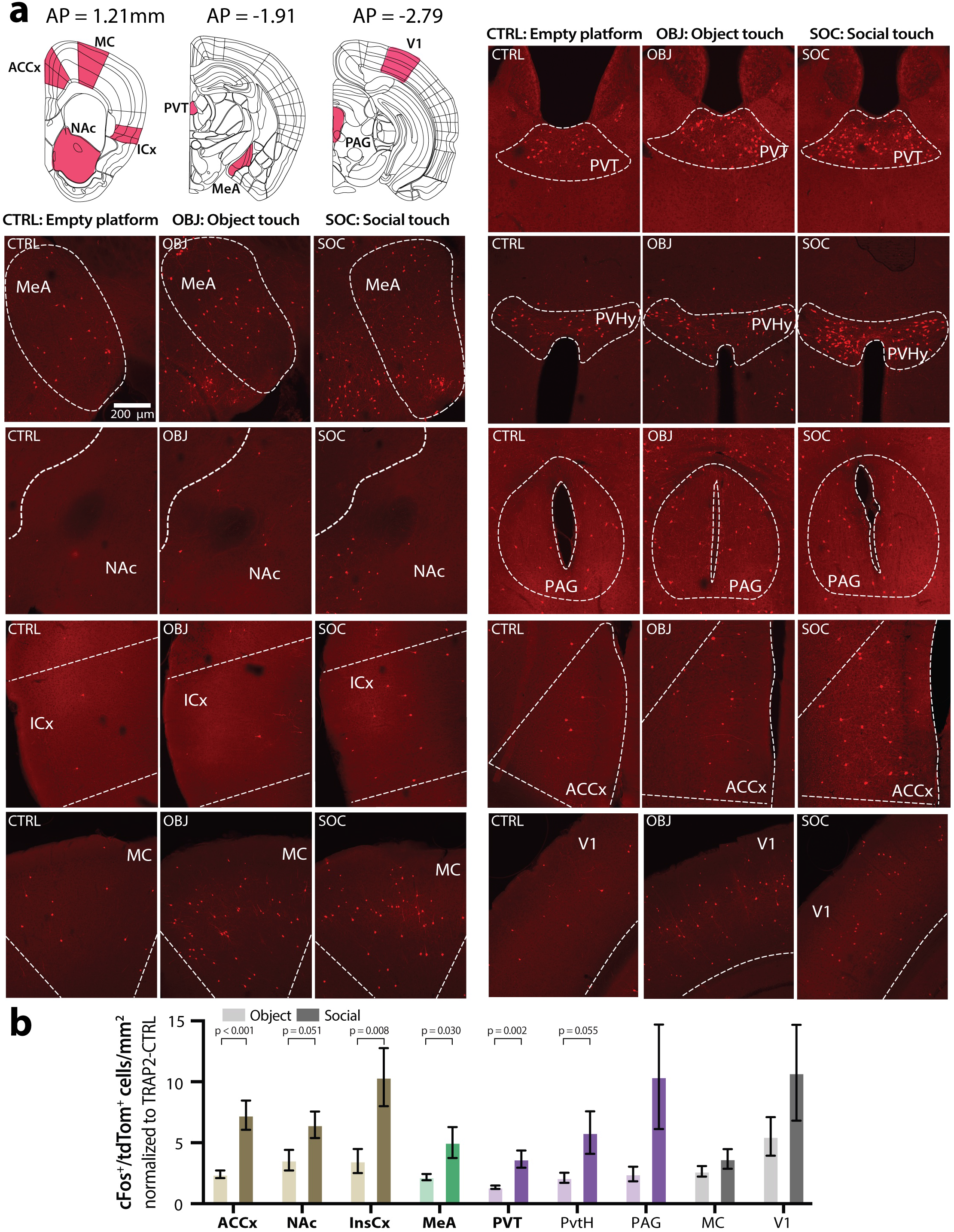
Differential cFos expression to object touch vs. social touch across relevant brain regions. **a.** Cartoons to indicate the locations across the anterior-posterior (AP) axis of the mouse brain where images of cFos expression in TRAP2 mice were taken (top left). Example images of cFos expression from medial amygdala (MeA), nucleus accumbens (NAc), insular cortex (ICx), motor cortex (MC), paraventricular nucleus of the thalamus (PVT), paraventricular nucleus of the hypothalamus (PVHy), periaqueductal gray (PAG), anterior cinculate cortex (ACCx), and primary visual cortex (V1) during object touch (OBJ) and social touch (SOC), compared to the no touch (empty platform) control (CTRL). **b.** Density of cFos-expressing (tdTom+) cells per mm^2^ for mice that received repeated presentations of OBJ or SOC touch, normalized to the CTRL cell density for each brain region (bottom). ***p<0.001, **p<0.01, *p<0.05, normality was tested with D’Agostino & Pearson test followed by unpaired nonparametric Mann-Whitney or parametric t-test for each brain region. Each bar represents data from 5-6 mice, and at least 6 images were collected from a single mouse for each brain region.

**Supplementary Fig. 3:**
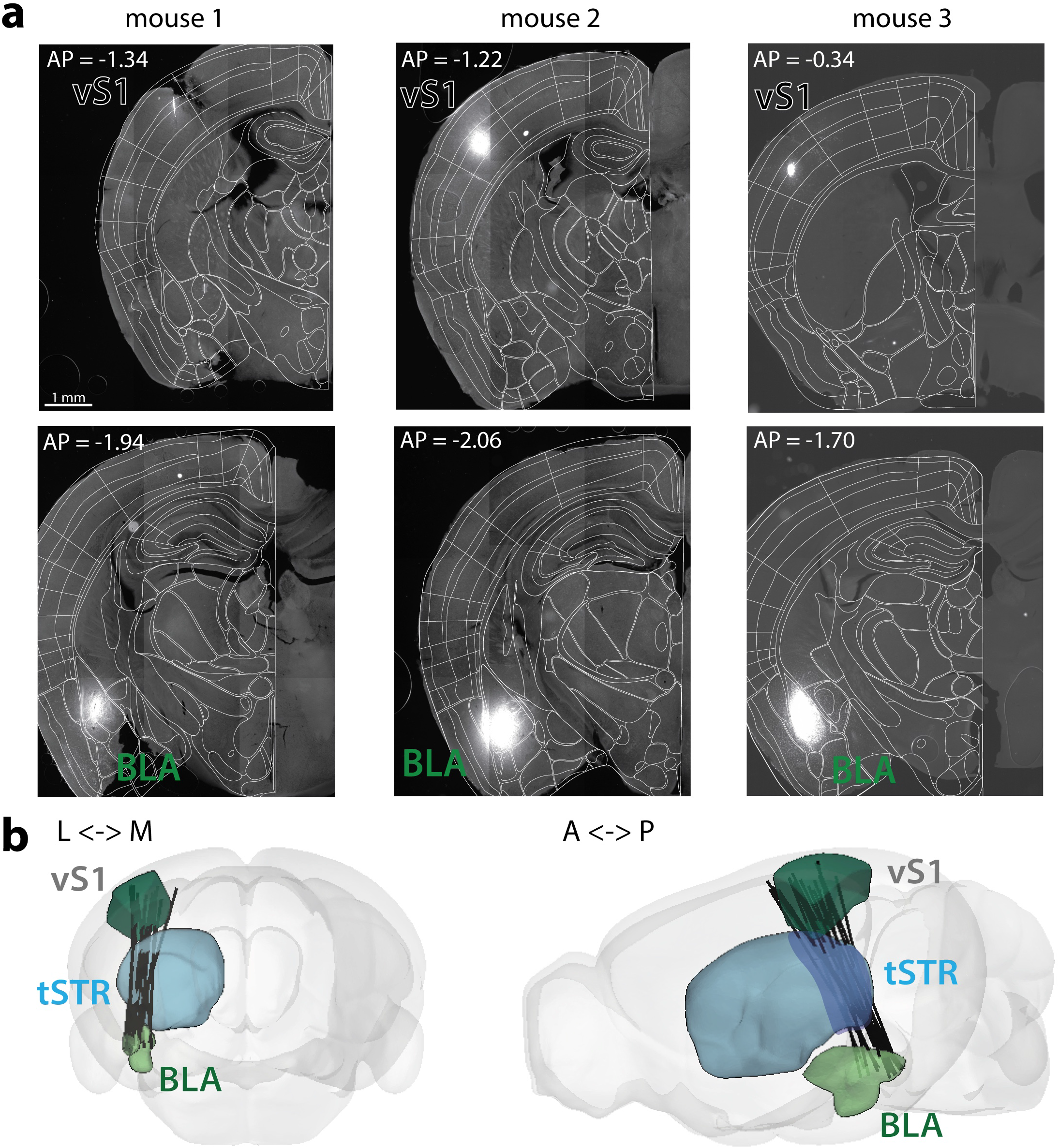
Histological reconstruction of the trajectory of Neuropixels probes. **a.** Example fluorescence imaging in three mouse brains of the probe shank trajectory (stained with DiI) to confirm accurate targeting of vS1, tSTR and BLA. Brain sections (50 µm thickness) are aligned to the Allen Mouse Brain Atlas (scale bar = 1 mm). **b.** 3D reconstruction views of Neuropixels probe tract across the brains of all WT (n=9) and *Fmr1* KO mice (n=10) using the location of DiI fluorescence in coronal brain slices. We confirmed that each probe traverses vS1, tSTR and BLA across the mediolateral (M-L) and anteroposterior (A-P) axes.

**Supplementary Fig. 4:**
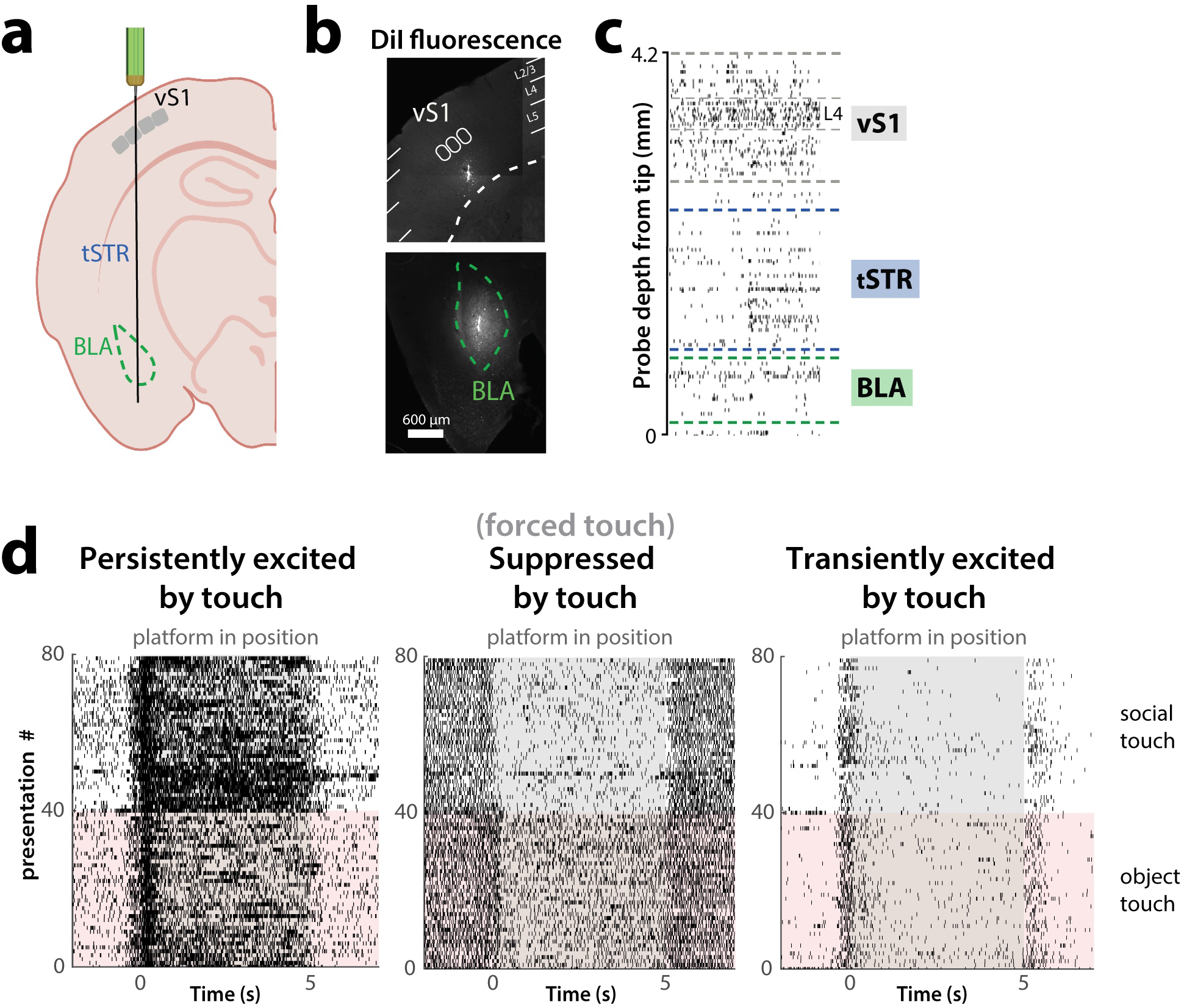
Approach to allocate individual units to vS1, tSTR or BLA. **a.** Cartoon showing Neuropixels probe implanted at 0° angle using mouse brain coordinates - 1.46 AP, 2.9 ML 3.75 DV to target vS1, tSTR and BLA. **b.** DiI fluorescence used to confirm probe targeting in vS1 and BLA (scale bar=600 µm). **c.** Patterns of action potential spiking across time and depth (in mm) were used to allocate units recorded from Neuropixels to different brain regions, in addition to the estimated depth of the unit inferred from the probe trajectory reconstructions. **d.** Example rasters of all action potential spikes across all 40 forced presentations of social touch (top) and object touch (bottom) for three example units that are persistently excited by touch, suppressed by touch, or transiently excited by the initial contact just before platform stops. Time 0 s denotes when the platform stops moving towards the test mouse and 5 s denotes when the platform starts moving away from the test mouse. Note how the left-most unit is preferentially excited by social touch, while the middle unit is preferentially suppressed by social touch.

**Supplementary Fig. 5:**
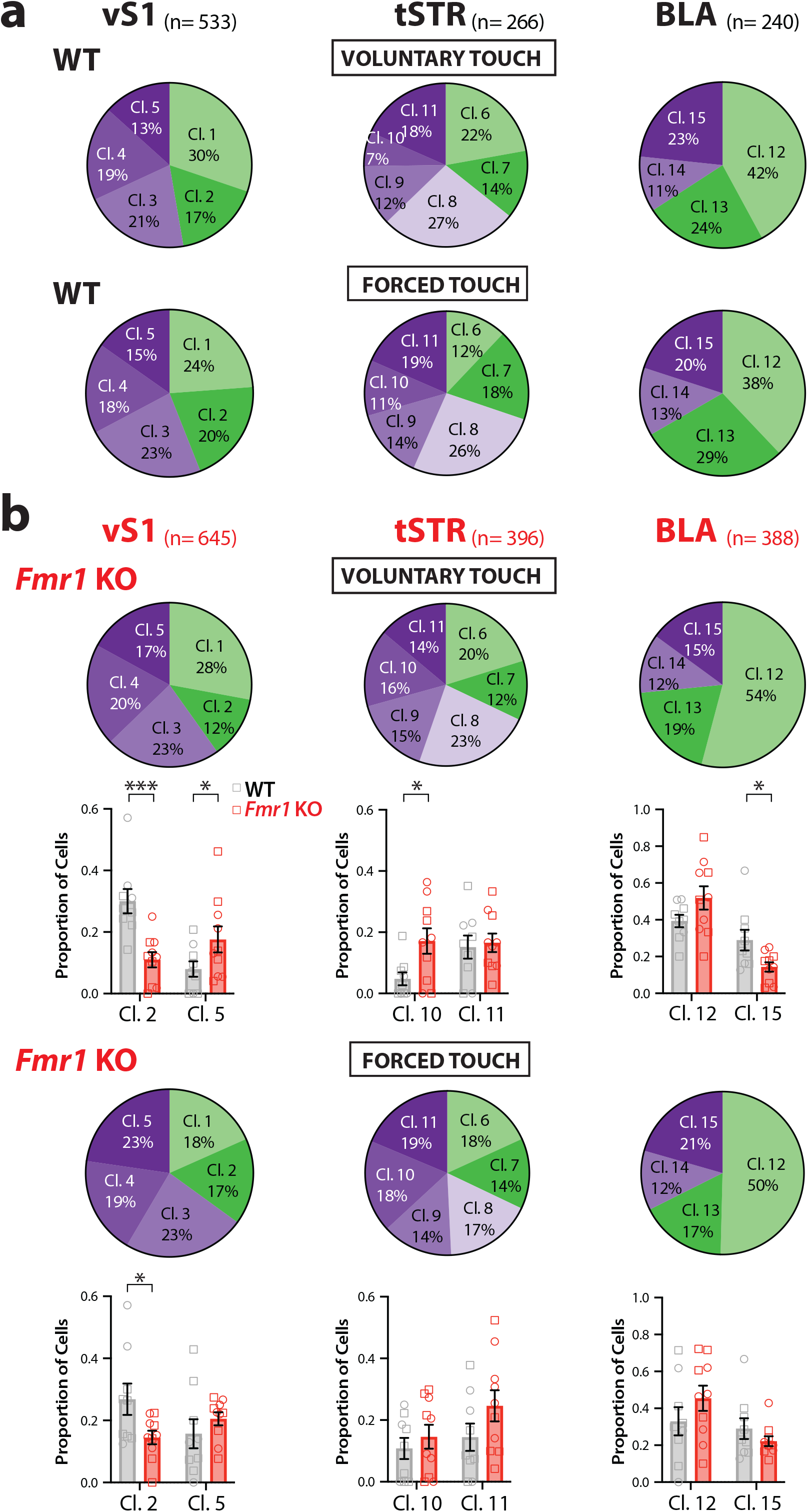
Proportion of total cells in each cluster differs within WT and between WT and *Fmr1* KO mice. **a.** Percentage of cells belonging to different clusters in vS1, tSTR and BLA for all WT mice (n=9) during voluntary touch (top) or forced touch (bottom). **b.** Percentage of cells belonging to different clusters for all *Fmr1*KO mice (n=10) (top). Quantification of the relative abundance of cells in various clusters (as a proportion of total cells) for different WT and *Fmr1* KO mice during voluntary and forced touch (bottom). ***p<0.001, *p<0.05 for unpaired parametric t-test WT vs *Fmr1* KO for each cluster. Squares=males, circles=females.

**Supplementary Fig. 6:**
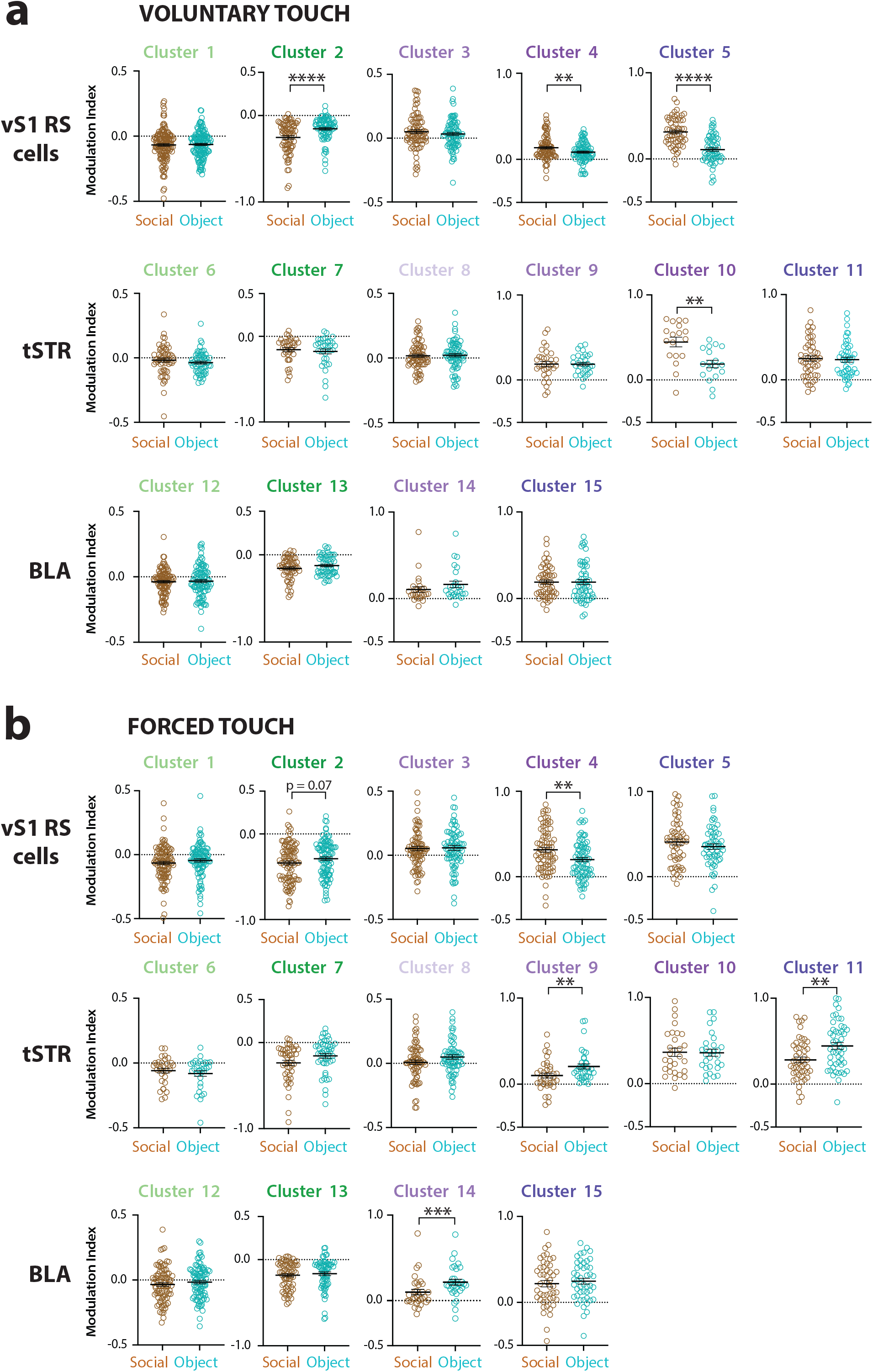
Clusters in vS1, tSTR and BLA are modulated differently by voluntary social and object touch. **a.** Modulation index of vS1 RS, tSTR and BLA cells in different clusters for social vs. object touch, during voluntary presentations, as an average of all 40 presentations. **b.** Same as in panel *a*, but for forced touch. ****p<0.001, *** p<0.005, **p<0.01 for paired parametric t-test. Each symbol represents a single cell taken from across 9 WT mice.

**Supplementary Fig. 7:**
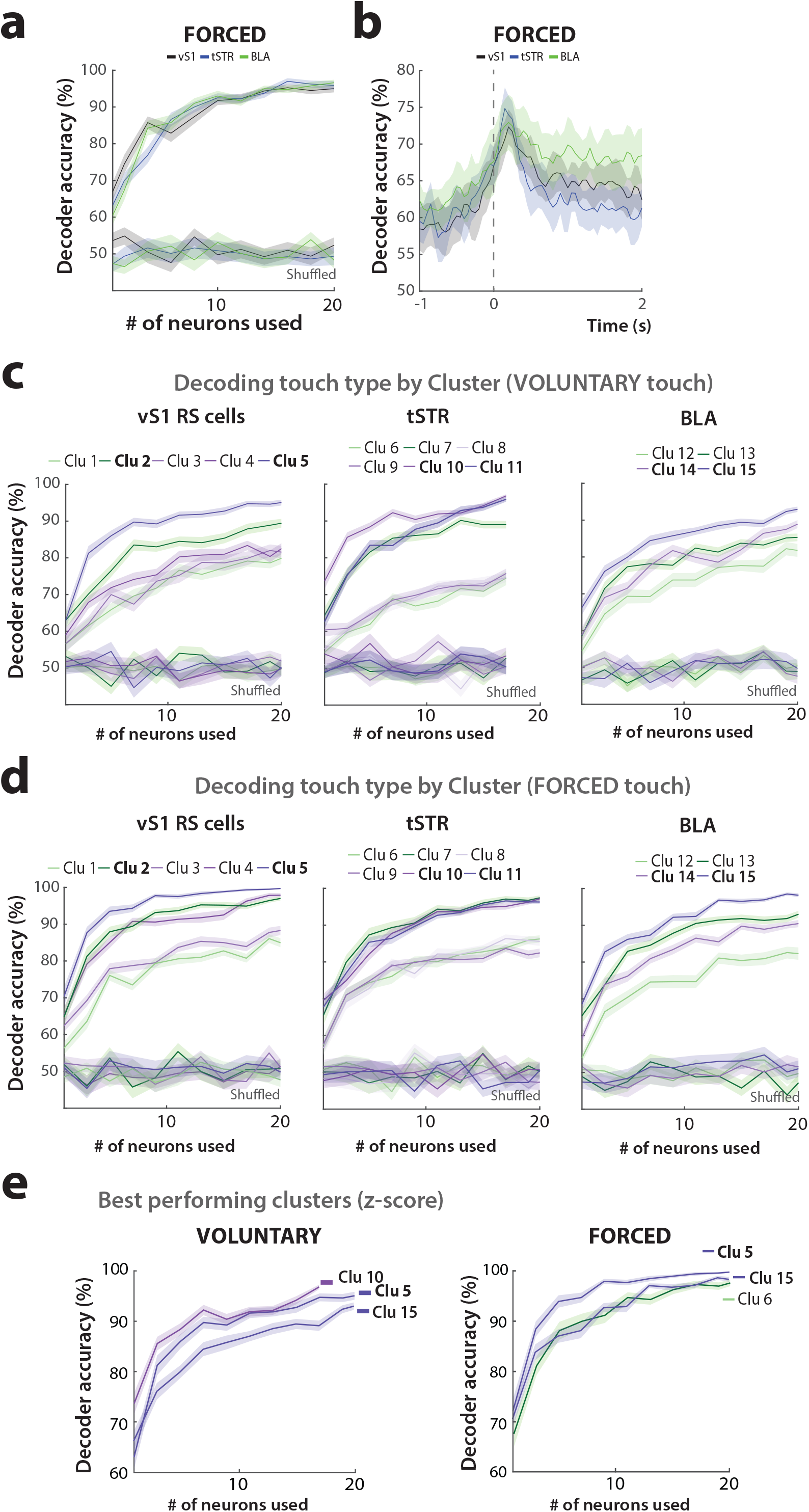
Decoding touch context (social vs. object) from mean neural activity in vS1, tSTR and BLA. **a.** Decoder accuracy for touch context based on the activity of cells in vS1, tSTR and BLA during forced touch (0 to 5 s). From each brain region, 1-20 neurons were randomly selected to be used in the SVM classifier. Decoder accuracy is also shown for shuffled data, where context identity was shuffled in 80% of object and social touch presentations (64 stimulations total) used for the training data set. **b.** Decoder accuracy for touch context based on activity of 10 randomly selected cells from each mouse in vS1, tSTR and BLA for every 50 ms during the stimulation period of forced touch (−2, 7s). **c.** Decoder accuracy as in panel *a* but for individual clusters in vS1 RS, tSTR and BLA, for voluntary touch. Decoder accuracy for tSTR clusters only goes up to 17 neurons because one tSTR cluster only had a total of 17 cells across all WT mice. **d.** Decoder accuracy as panel *c* but for forced touch. **e.** Decoder accuracy just as in panels *c-d*, but using activity of the best performing clusters in vS1 RS, tSTR and BLA for voluntary (left) and forced presentations (right).

**Supplementary Fig. 8:**
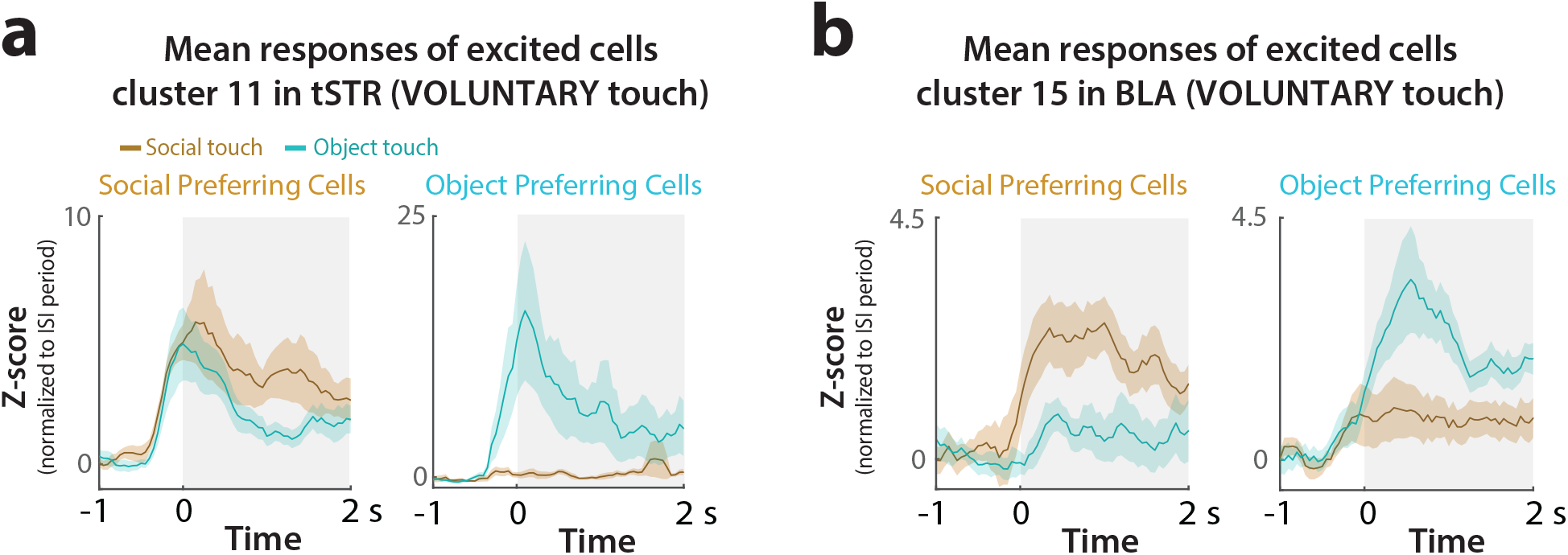
Responses of social-preferring and object-preferring cells in tSTR and BLA. **a.** Averaged z-score firing rate of all social and object preferring cells in Cl. 11 of the tSTR during voluntary object and social touch. **b.** Just as in panel *a* but for cells in Cl. 15 of the BLA.

**Supplementary Fig. 9:**
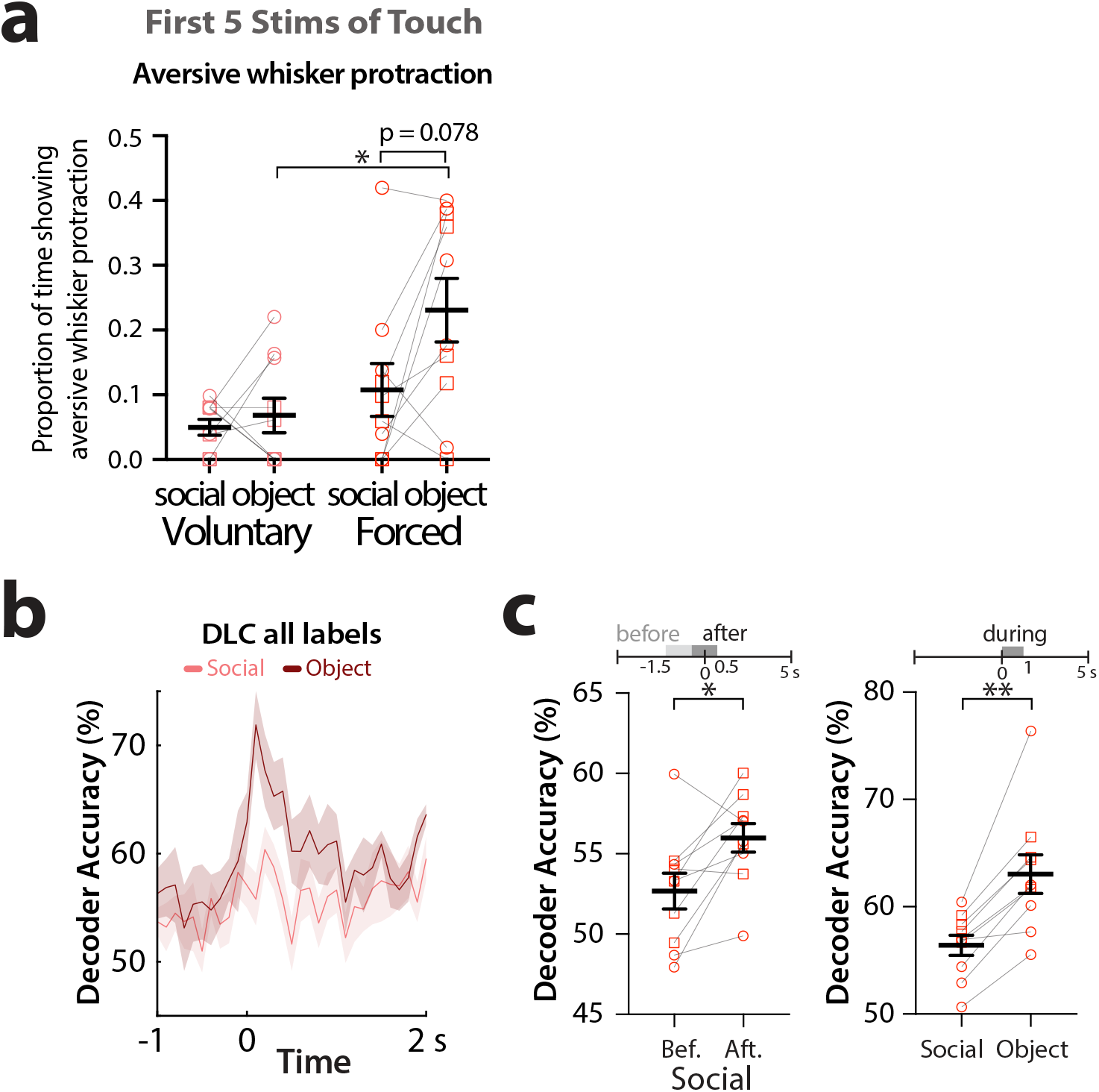
*Fmr1* KO mice show similar levels of aversive whisker protraction for social touch and object touch and find social touch within their personal space more aversive. **a.** Proportion of time that *Fmr1* KO mice exhibit sustained aversive whisker protraction during social touch and object touch for voluntary and forced presentations. Squares=males, circles=females. *p<0.05 for two-way ANOVA with Bonferroni’s. **b.** Decoder performance for touch choice (voluntary vs. forced) discrimination in *Fmr1* KO mice using SVM classifiers trained on all DeepLabCut (DLC) labels on the mouse’s face across time for social touch and object touch (from −1 s to +2 s after platform stops). **c.** Decoder accuracy for touch choice discrimination in *Fmr1* KO mice using all DLC labels before (−1.5 to −0.5 s) and after (−0.5 to 0.5 s) platform stops (left), or during the first second after platform stops for social touch and object touch (right). *p<0.05, **p<0.01 for parametric paired t-test for both panels.

**Supplementary Fig. 10:**
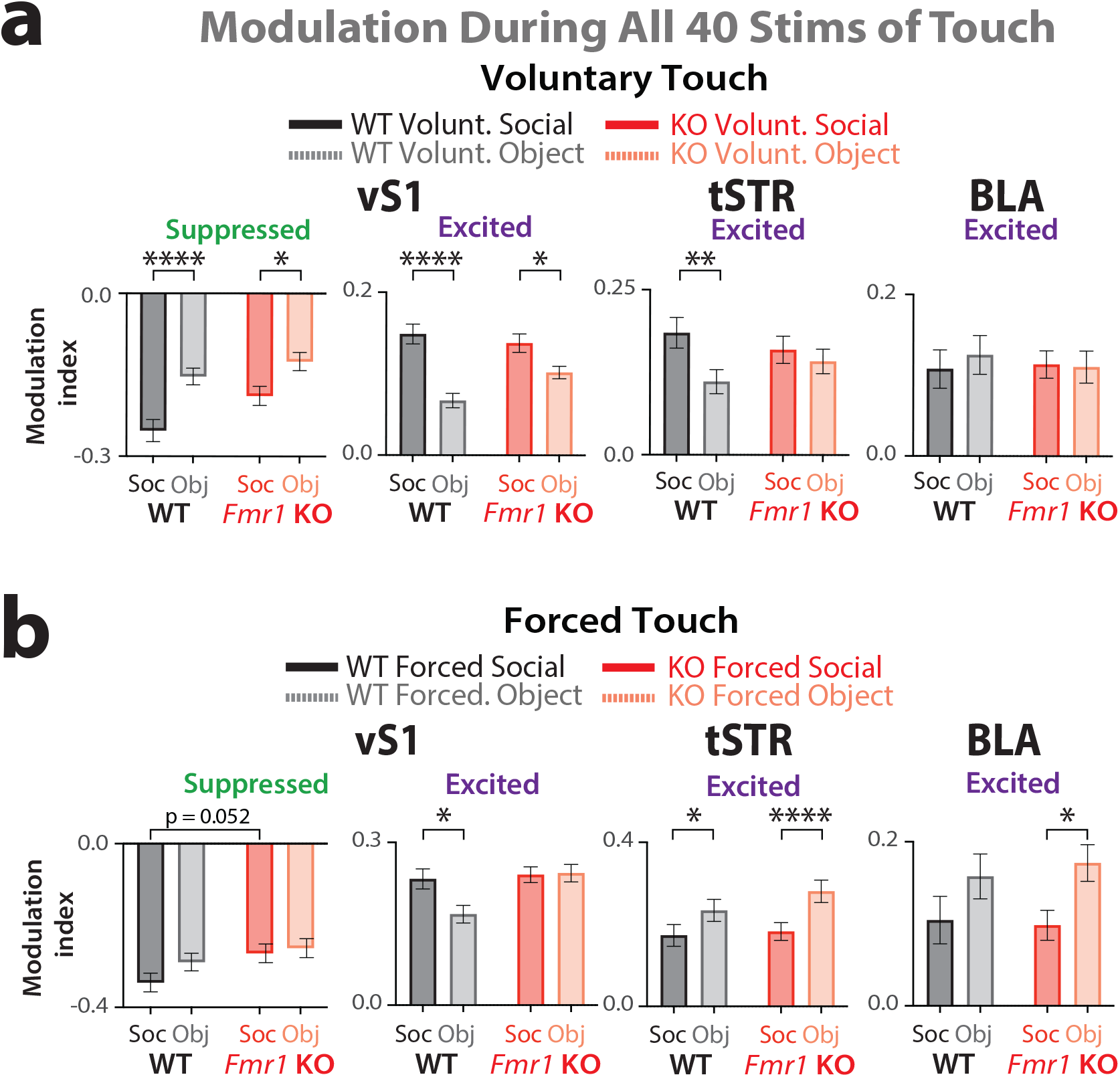
Modulation indices of vS1, tSTR and BLA cells in WT versus *Fmr1* KO mice for social and object touch. **a.** Modulation index of vS1 RS cells excited and suppressed by touch, as well as excited cells in tSTR and BLA in response to voluntary social vs. object touch, as an average of all 40 stimulations, in WT and *Fmr1* KO mice (n=9 and 10 respectively). ****p<0.001, **p<0.01, *p<0.05 for two-way ANOVA with Bonferroni’s. **b.** Same as panel *a* but for forced touch presentations.

**Supplementary Fig. 11:**
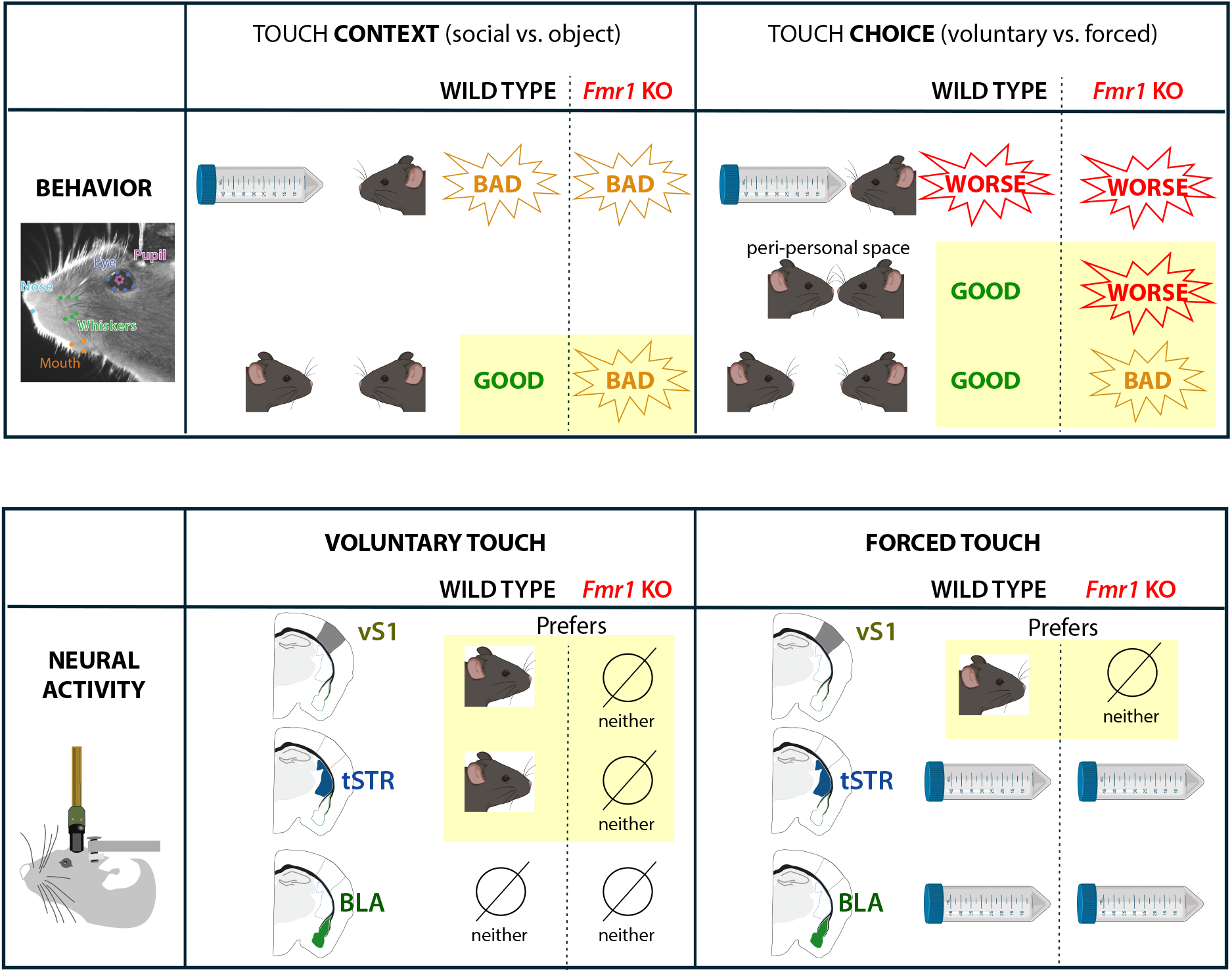
Summary of main findings. Here we summarize the main results related to behavioral responses (top) and neural activity (bottom). We compare how behaviors and neural activity differ according to differences in touch context (social versus object) and touch choice (voluntary versus forced interactions). For behavior, “good” means mice show minimal or no aversion; “bad” and “worse” means the mice experience increasing degrees of aversion. We also illustrate how response of *Fmr1* KO mice differ from those of WT mice (note that yellow highlights underline the differences in behavior and neural activity between genotypes).

